# Structure determination and dual targeting of a plant TACO1 identifies its ancient role as an organelle translation regulator

**DOI:** 10.64898/2026.05.28.728419

**Authors:** Parth K. Raval, Nuša Kos Thaler, Cristopher Mitchell, Maria Lozano-Quiles, Tommi Kajander, Daniel W. Djamriani, Jens Reiners, Sander HJ Smits, Sarah J. Butcher, Brendan J. Battersby, Sven B. Gould

## Abstract

Ribosome stalling caused by polyproline (PPs) motifs is common. Their translation is enhanced by accessory proteins such as YebC in bacteria, whose homolog, TRANSLATIONAL ACTIVATOR OF CYTOCHROME C OXIDASE 1 (TACO1), aids the translation of mitochondria-encoded proteins. The prevalence of PP motifs across plastid-encoded genes and their impact on the translation of photosynthesis-relevant proteins remains unexplored. Equally, a translation-enhancer of PP motifs equivalent to TACO1 for plastid ribosomes has not been reported. Here, we show that plastid genomes encode 24 proteins with a minimum of one PP motif on average, half of which are conserved in their cyanobacterial homologs. We also find that the vast majority of eukaryotes, including plants, encode only a single TACO1 that we demonstrate to be dually targeted to mitochondria and plastids of *Marchantia polymorpha.* We resolved the MpTACO1 structure at 2.34 Å by X-ray crystallography and its flexibility by small-angle X-ray scattering. Through modelling, we demonstrate that MpTACO1 can fit into the peptidyl transfer centre of plant chlororibosomes in a similar manner as human TACO1 in the mitoribosome. The identification and structure determination of the first plastid-targeted YebC/TACO1 allows us to sketch a unified model for the function and evolution of this ancient family of ribosomal accessory proteins, underscoring their indispensable role in the translation of bioenergetic membrane proteins reaching back almost 4 billion years.

**Highlights:**

- Dozens of GC-rich polyproline (PP) encoding regions are retained by AT-rich genomes
- PP motif conservation hints at regulatory mechanisms and required translation pauses
- Chloroplast targeting of a (mitochondrial) translation enhancer of PP motifs
- MpTACO1 structure at 2.34 Å resolution demonstrates its high level of conservation

## Introduction

Plant cells are the product of serial endosymbiosis. The timing of the events remains uncertain, but occurred in the Palaeoproterozoic era^1^ and first gave rise to eukaryotes through the origin of mitochondria, and subsequently to phototrophic eukaryotes through the origin of the plastid^2–6^. The transition from a bacterial endosymbiont to a eukaryotic organelle entails the transfer of bacterial genes to the nucleus through endosymbiotic gene transfer (EGT)^7–10^. Thousands of organellar proteins are hence translated in the cytosol and must be imported by the organelles of endosymbiotic origin^11–14^. Despite EGT and the emergence of dedicated protein import systems, both mitochondria and plastids still encode a small, albeit key, subset of genes that are translated into proteins in their matrix and stroma, respectively.

For both organelle genomes, one can sort most of their genes into those encoding either the remaining subunits of the ribosomes or proteins of the oxidative phosphorylation complexes of mitochondrial respiration and plastid photosynthesis^15–17^. Some ribosomal proteins remain organelle-encoded due to functional constraints associated with ribosomal complex assembly^16^, while bioenergetic membrane protein-encoding genes are likely retained due to issues associated with their hydrophobicity^18–21^ and *in organello* redox regulation^22–24^. Organelle-encoded complexes require nuclear-encoded subunits to function, some of which are dual targeted to both organelles after the translation from a single gene, e.g. rpL10 or rpS16^25,26^. The maturation of bioenergetic membrane complexes is hence a delicate process that requires a spatiotemporal orchestrated assembly of factors encoded by two genomes. The principal components and the core mechanism of protein translation and membrane insertion were inherited by eukaryotic organelles from their bacterial ancestors and remain well conserved^27,28^.

Ribosomes, whether cytosolic, mitochondrial or plastidial, can stall during translation elongation due to mRNA structures such as hairpins or challenging amino acid sequences, most notably proline residues^25,29–35^. Proline’s unique cyclic structure reduces conformational flexibility during the formation of peptide bonds, which can form trans- and cis-configurations at the peptidyl transferase center (PTC) of the ribosome, resulting in significantly slower translation rates^35,36^. Several accessory factors evolved early in evolution to aid the peptide bond formation and prevent stalling, specifically for the translation of two or more consecutive prolines (hereafter referred to as polyproline or PP motifs). These include the bacterial Elongation Factor P (EF-P), the archaeal Initiation Factor 5A (aIF-5A), members of the ABC-F ATPases, and YebC^37,38^. In mitochondria, a homolog of YebC called TRANSLATIONAL ACTIVATOR OF CYTOCHROME C OXIDASE 1 (TACO1) is required for the mitochondrial synthesis of COX1 and, potentially also COX3^39–42^. Human COX1 encodes no less than five PP motifs of which at least three have been found to stall ribosomes^39^. Recent *in organello* cryogenic electron microscopy (cryoEM) found TACO1 interacting with the mitochondrial 16S rRNA near the A-site, bridging the small and large mitoribosomal subunits^43,44^. YebC enhances the translation of polyproline stretches in bacteria^37^ and was recently shown to transiently interact with the bacterial 23S rRNA of the 70S ribosome^38^. A model is emerging whereby factors such as YebC and its eukaryotic homolog TACO1 stabilize the A-site prolyl-tRNA during peptide bond formation on bacterial and mitochondrial ribosomes^43,44^.

Plastids are of cyanobacterial origin and like most bacteria, cyanobacteria encode a YebC homolog (Fig. 1, Fig. S1B). There are, however, neither reports of the cyanobacterial YebC’s fate in algae and plants, nor the prevalence and potential impact of prolines and PP motifs on protein translation in plastids. This presents us with an unexplored phenomenon of plastid biology. Here, we screened 15,208 algae and plant plastid genomes and found that, on average, every genome encodes a quarter of the genes with at least one PP motif. Of the 169 nuclear genomes of photosynthetic and non-photosynthetic eukaryotes we screened, 122 encode only a single YebC/TACO1 homolog, which we demonstrate is dual targeted to mitochondria and plastids of the liverwort *Marchantia polymorpha*. We solved the crystal structure of *M. polymorpha* TACO1 (MpTACO1), revealing a highly conserved structure, and further using SAXS revealed three conformations in solution. Our data suggest that the plant YebC/TACO1 homolog is a conserved protein critical for the translation regulation of organelle-encoded proteins not only in mitochondria, but also in the plastids of algae and plants.

**Fig. 1:**
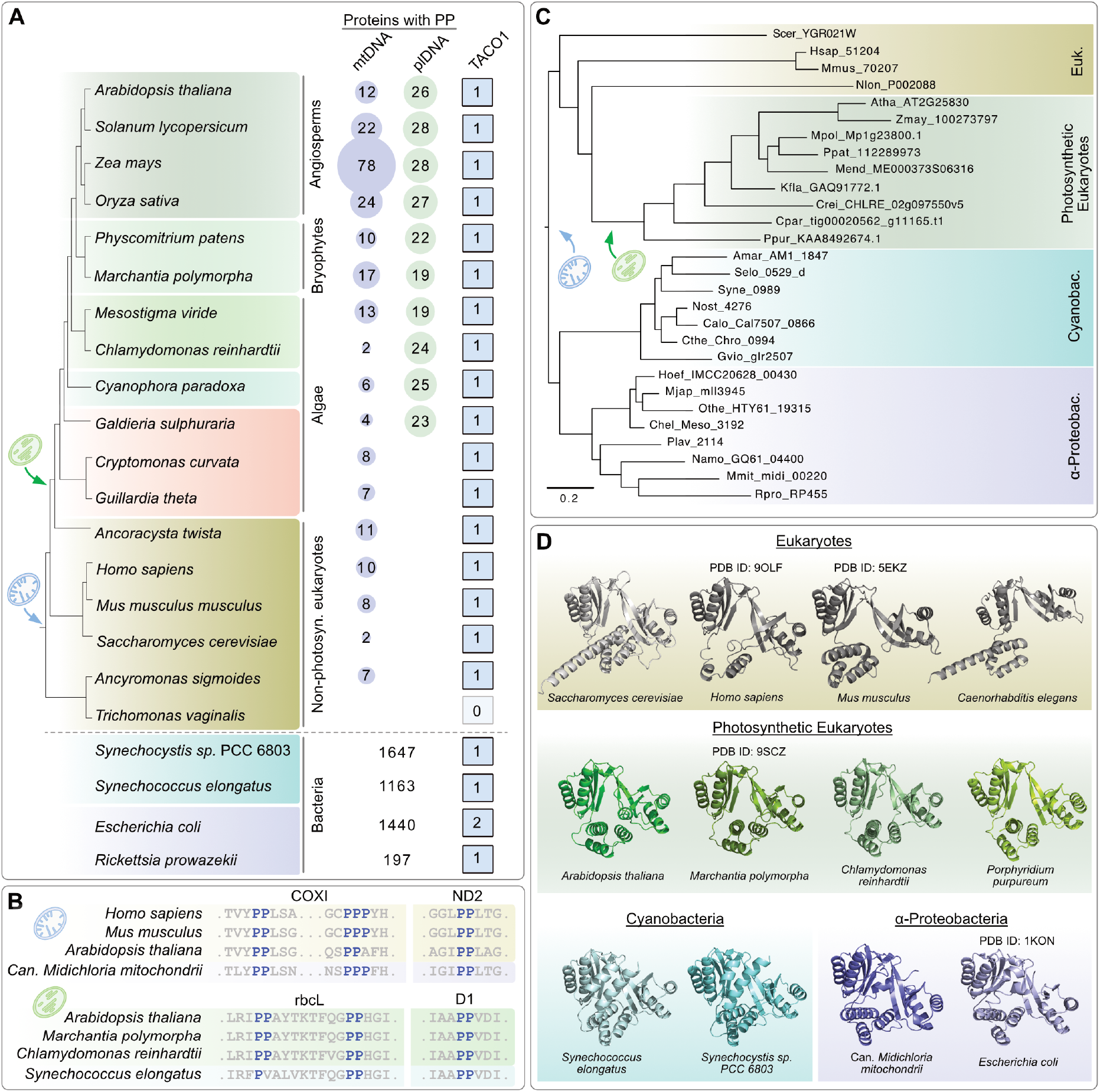
Conservation of YebC/TACO1 homologs and PP motifs across bacteria and eukaryotes. **(A)** Number of polyproline (PP) motifs containing proteins in bacterial, mitochondrial and plastid genomes along with the number of genome-encoded TACO1 homologs across diverse eukaryotes and prokaryotes. **(B)** Examples of PP motifs that are present in alpha-proteobacteria and cyanobacteria, and which were inherited by eukaryotes through endosymbiosis and remain organelle-encoded. **(C)** Phylogenetic tree of YebC/TACO1 genes, rooted between prokaryotes and eukaryotes. **(D)** Experimentally resolved (human PDB ID:9OLF, mouse PDB ID:5EKZ, *E. coli* PDB ID:1KON, and in this study *M. polymorpha* PDB ID:9SCZ) and predicted AlphaFold3^48^ structures of YebC/TACO1 proteins from selected diverse eukaryotes and prokaryotes.

## Results

### Organelle-encoded PP motifs and YebC/TACO1 proteins share an ancient history

While PP motifs are sporadically discussed for individual proteins or a species group, their global frequency and the level of conservation across distant taxa has not been explored, especially not for plastid genomes. We screened 18,059 mitochondrial and 15,208 plastid genomes of diverse eukaryotes. Mitochondrial genomes encode six proteins with at least one PP motif on average (42.6% of the organelle-encoded genes), while plastid genomes encode 24 (29.4%) (Fig. 1A, Fig. S1A). The positions of many PP motifs are evolutionarily conserved in bacteria, animals and plants (Fig. 1B, Fig. S2-3) and includes motifs of COXI and the photosystem II protein D1 (psbA) that are found in the vicinity of transmembrane domains. As prolines are encoded by the CCN codon, this speaks for a >1.5 billion years long retention of these GC-rich stretches by plastid genomes that evolve under evolutionary pressures that favour AT-rich genomes ^45,46^. For comparison, 56 diverse alpha-proteobacterial and 63 cyanobacterial genomes encode about a thousand PP-containing proteins each (Fig. 1A, Fig. S1A).

The search for YebC/TACO1 homologs among the genomes of 113 bacteria and 169 eukaryotes finds that 107 (94.6%) of the prokaryotes and 122 (72.1%) of the eukaryotes encode only a single copy (Fig. 1A, Fig. S1B). Eukaryotic species with more than one copy include a small subset of plants with extreme polyploidy and early-branching eukaryotes, for which their draft genomes returned only partial YebC/TACO1-like sequences (Fig. S1A). In any case, a majority of land plant species with extreme ploidy and those that are otherwise characterised by an expansion of organelle-targeted proteins^47^, encode only a single YebC/TACO1 protein. Eukaryotes lacking YebC/TACO1 homologs include parasites such as *Giardia intestinalis* and *Trichomonas vaginalis,* whose mitochondria-derived organelles lack genomes and hence do not translate proteins to begin with, underscoring YebC/TACO1’s role as an accessory protein of mitoribosomes which trace their ancestry to bacterial ribosomes.

A phylogenetic analysis places all prokaryotic YebC proteins from both alphaproteobacteria and cyanobacteria – representing the donor groups to the endosymbiotic origin of mitochondria and plastids, respectively – as a monophyletic sister clade to eukaryotic TACO1 sequences, where algae and plant TACO1 homologs form a monophyletic group (Fig.1C). Due to the high level of sequence conservation, a definite claim on whether the alphaproteobacterial (mitochondrial) or cyanobacterial (plastid) YebC has been retained in plants is not possible. The sequence conservation is echoed by the predicted and experimentally validated structures of YebC/TACO1 proteins from across the tree of life (Fig. 1C-D, Fig.2, S4A). Structure-based dendrograms and clustering also place prokaryotic homologs together as a monophyletic sister clade to eukaryotes (Fig. S4B-D). Taken together, our analyses underscore an ancient and abundant occurrence of PP motifs across prokaryotic genomes, some of which remain encoded by the two organelles of endosymbiotic origin, and that a single TACO1 protein associated with their translation, at least in mitochondria, is present in the vast majority of eukaryotes, including algae and plants.

**Fig. 2:**
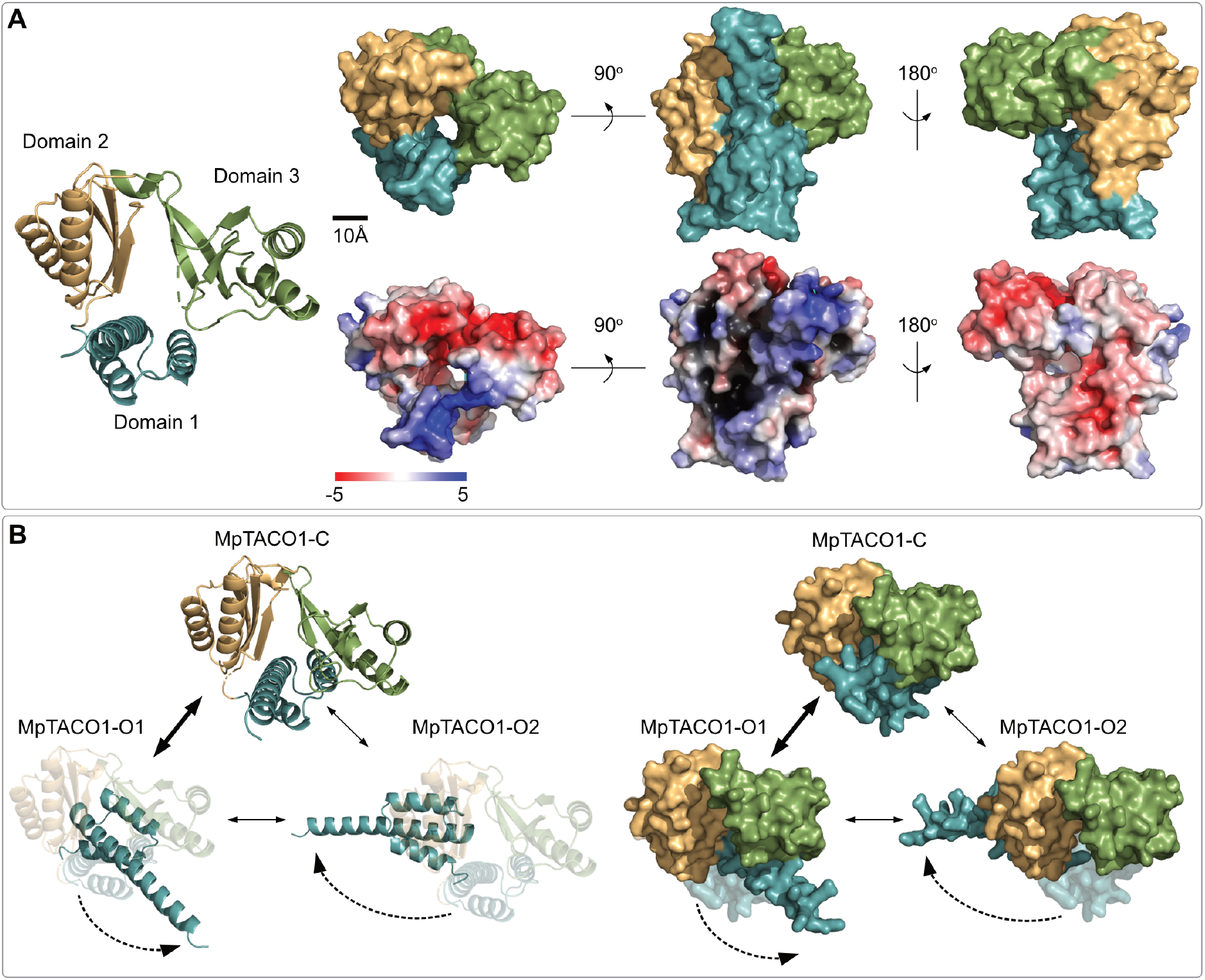
MpTACO1 contains a conserved hinge and forms three distinct conformations in solution. **(A)** Crystal structure of MpTACO1 represented as a ribbon-based cartoon, along with surface views colour-coded by domains (top) and by local electrostatic potentials (bottom). **(B)** Three conformations present within the ensemble at 40%, 50% and 10%, labelled MpTACO-Closed (MpTACO-C), MpTACO-Open1 (MpTACO-O1) and MpTACO-Open2 (MpTACO-O2). MpTACO-C resembles the most compact structure solved from X-ray crystallography and is shown as a semi-transparent structure underneath O1 and O2 conformations to indicate the relative motion of domain 1. Arrow thickness indicates the dominance with which the conformational change was measured.

### X-ray crystallography and SAXS uncovers three distinct conformations of MpTACO1

Thus far, the worldwide protein data bank houses five YebC/TACO1 family atomic models for bacteria and mammals, with no entries for plants. To fill this gap, we recombinantly expressed the Marchantia homolog MpTACO1 (Mp1g23800; coding for amino acids 127-374) in *E. coli.* Amino acids 1-126 were omitted, because a protein alignment of eukaryotic TACO1 and bacterial YebC sequences suggested this sequence to represent the N-terminal targeting peptide (TP; Fig. S5A). TPs tend to be unstructured, can interfere with crystallisation and offer no information on the function of a mature protein. MpTACO1 lacking the TP was purified as a monomer, crystallized, and we determined its structure with X-ray diffraction using molecular replacement (Fig. S5B-F). We obtained a resolution of 2.34 Å with 97.61% completeness, which allowed for the atomic modelling of residues 127-293 and 301-373, comprising the fully encoded construct except for 7 residues in a flexible loop region of domain 3, with no Ramachandran or rotamer outliers, (Fig. S5, Table S1). MpTACO1 comprises three domains (Fig. 2A): domain 1 (residues 127-200,) consists of a bundle of three α-helices, domain 2 (residues 201-253, 339-373) features three antiparallel β-strands facing two α-helices, and domain 3 (residues 254-338) contains four antiparallel β-strands that face two α-helices (Fig. 2A). The surface of domain 1 is primarily positively-charged, domain 2 has mixed charges, and domain 3 is primarily negatively-charged (Fig. 2A).

The structure of TACO1 homologs has been reported to exhibit both an open and closed cleft between domains 2 and 3, previously thought to be an RNA-binding cleft, suggesting that conformational changes are possible^40,49,50^. To analyse the behaviour of MpTACO1 in solution, we performed small- angle X-ray scattering (SAXS) experiments. The SAXS results also showed that MpTACO1 is a monomer in solution (see Table S2). The crystal structure of MpTACO1 agreed with the experimental SAXS data (χ² value 1.39), however, a better fit was achieved by ensemble modelling (Fig. 2B). Based on the Kratky plot, MpTACO1 is a compact molecule with a certain degree of flexibility, which is consistent with the presence of open and closed conformations inferred from previous studies^40,49,50^ (Fig. S6). To analyse this movement in solution, we used an Ensemble Optimisation Method (EOM). In this analysis, domain 1 is free to move in solution and the resulting *R_g_* distribution primarily exhibits two volume fractions with *R_g_* values of 2.01 nm and 2.11 nm, respectively (Fig. 2B, Fig. S6). The protein maintains an overall compact conformation, representing the closed (MpTACO-C, volume fraction 40%) and open conformations (MpTACO-O1, volume fraction 50% and MpTACO-O2, volume fraction 10%) within the ensemble (Fig. 2B, Fig. S6), with the closed conformation being similar to that of the crystal structure and to that of the human TACO1 (HsTACO1) modelled in the bound state to human mitoribosomes^43^. Utilizing the published structure of HsTACO1 in the human mitoribosome (PDB ID 9OLF,^43^) and the MpTACO1 crystal structure, we sought to determine if MpTACO1 could interact in a similar position in the chlororibosome. Domain 3 of HsTACO1 has an extended interaction interface with ribosomal protein bL12 (Fig. 3a,c). The residues that comprise this interface exhibit a high degree of conservation in relation to residues outside of the interface (Fig. 3b). After identifying homologous interacting residues in the MpTACO1 crystal structure and predicted *M. polymorpha* bL12 structure, we were able to dock the subunits, simulating an interaction interface that is similar to the known human interface (Fig. 3c,d). There are eleven residues in the interface that have conserved sidechain properties and only two differences that alter the polarity or charge of the sidechains between the human interface and the simulated *M. polymorpha* interface, one in bL12 (V159 to K306) and one in TACO1 (E208 to L151).

**Fig. 3:**
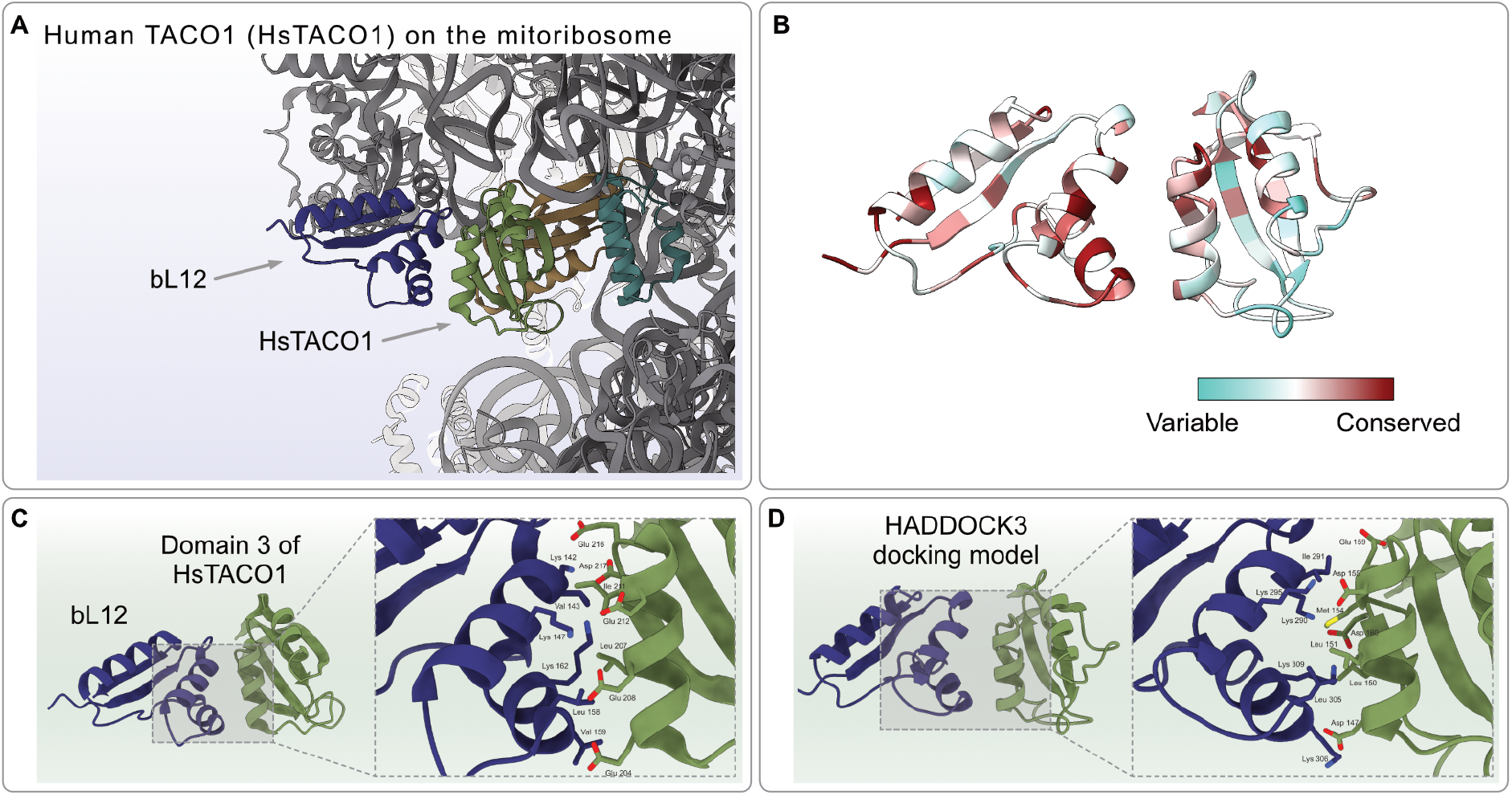
Modelling of TACO1 interactions with the entry tunnel of the ribosome. **(A)** The human mitoribosome structure (PDB ID 9OLF^43^) is depicted in a grey cartoon representation with bL12 highlighted in dark blue. The three domains of the HsTACO1 are coloured as in Fig. 2. **(B)** Domain 3 of the *M. polymorpha* TACO1 (PDB ID 9SCZ) crystal structure and AlphaFold3 bL12 models coloured by residue based on conservation, as calculated by ConSurf^51,52^, depicted as a gradient from turquoise (variable) to red (conserved). **(C)** Interface between domain 3 of HsTACO1 and bL12 (PDB ID 9OLF), with a close-up showing potential interacting sidechains in stick representation, coloured by heteroatoms (nitrogen: blue, oxygen: red, sulphur: yellow). **(D)** Predicted interface between domain 3 of the *M. polymorpha* TACO1 crystal structure (PDB ID 9SCZ) and an AlphaFold3-generated bL12 model, created through HADDOCK3 docking simulations^53^. Homologous residues to those shown in (C) are shown in a close-up with their sidechains in stick representation, coloured by heteroatoms.

### MpTACO1 is dual targeted to mitochondria and plastids

We fused the full-length sequence of MpTACO1 to citrine and determined its localization *in vivo* in *M. polymorpha*. The construct was targeted to mitochondria and plastids and the citrine signal appeared to be equally distributed between both compartments in the gemmae and adult thalli stages of the liverwort (Fig. 4A, Fig. S7A. Fig. S8). As with the full-length sequence, the first 105 amino acids of MpTACO1 were sufficient to sort the reporter to both mitochondria and plastids in protoplasts, which also allow for a better visualization of stained mitochondria (Fig. 4B, Fig. S7B, Fig. S9). Finally, to substantiate the plastid localization of MpTACO1 independent of microscopy, we isolated plastids from two independent transfectants and validated the presence of the full-length construct through immunoblotting (Fig. S10).

**Fig. 4:**
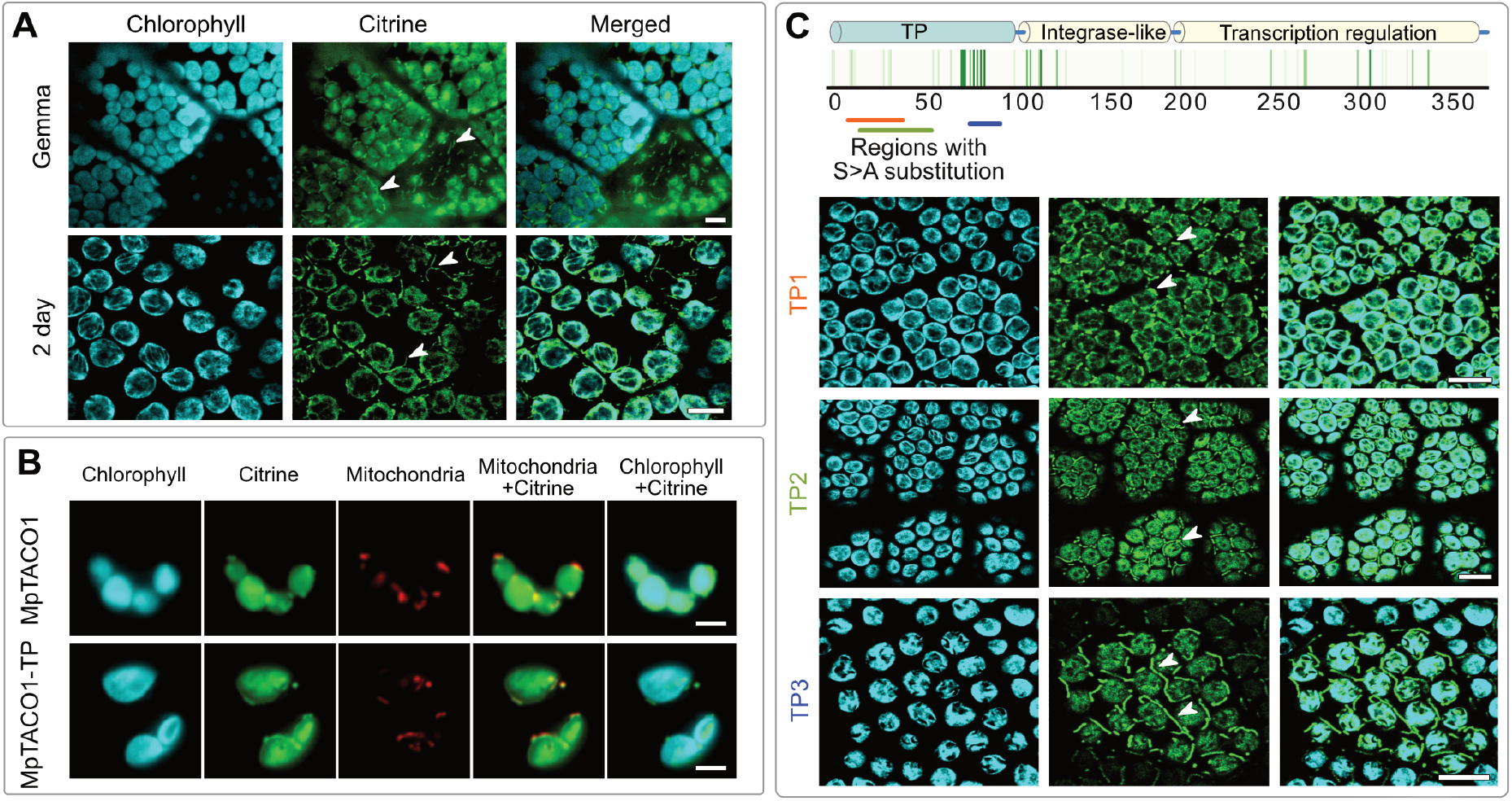
Protein dual targeting of TACO1 in the liverwort *Marchantia*. **(A)** Intracellular localisation of MpTACO1 in gemmae and two-day old thalli of *M. polymorpha.* **(B)** Images of protoplasts generated from thalli of one week old isogenic plant lines expressing both full MpTACO1 and only its targeting peptide (TP) fused to citrine. **(C)** Likelihood of serine phosphorylation in the TP of MpTACO1 depicted by shades of green, where each bar represents an amino acid residue. Key serine residues were mutated to alanine in three constructs: TP1 shown in orange (substituted residues: 4, 5, 14-17, 33, 36, 37), TP2 in green (substituted residues: 4, 5, 14-17, 33, 36, 37, 52, 54, 55), and TP3 in blue (substituted residues: 77-79, 82, 84, 86). Average phosphorylation probabilities of the mutated serine motifs were 0.17 (TP1), 0.32 (TP2) and 0.71 (TP3). Plastids were imaged through their autofluorescence and protoplast mitochondria through MitoTracker Red. Reticulate mitochondria are indicated by white arrow heads. Scalebars 10µm. Plastid autofluorescence during Citrine readout was filtered out while taking images for A and C, on a Leica TCS SP8 platform (Wetzlar, Germany) with time gating^56^, substantiating plastid localisation of this hitherto mitochondrial protein.

Plastid targeting peptides (pTPs) are usually enriched in amino acids that can be phosphorylated and the more N-terminal regions of mitochondrial TPs in positive charges^11,54^. Dual targeted proteins require a balance of both these features and indeed the TP of MpTACO1 encodes 22 serine residues with a mean phosphorylation probability of 0.45 and the first 25 amino acids have an overall charge of +4 (Fig. 4C). The abundance of serine residues suggests a degree of redundancy that may maintain and/or regulate dual targeting. To probe this, we substituted selected serine with alanine residues and generated three constructs, where 6-12 key serine residues were altered in total. Dual targeting to mitochondria and plastids was maintained by all three constructs, even upon the substitution of twelve serine residues as in construct TP2 (Fig. 4C, S7C). It suggests that MpTACO1 evolved a TP that can maintain dual targeting robustly, supporting its crucial role in both organelles. Such dual targeting of a single protein is not unusual in plants, especially for information-processing proteins^54,55^ and indeed, next to TACO1, we show that three additional proteins associated with transcription and translation are dual targeted in Marchantia (Fig. S11).

### Proximity labelling by MpTACO1 support a role in organelle protein biosynthesis

To identify potential interaction partners of MpTACO1 and further verify the dual localisation of the protein in the vicinity of organelle ribosomes, we fused the full-length protein to miniTurbo, a modified biotin ligase^57^. We selected for isogenic Marchantia plant lines expressing the construct in the absence of antibiotic selection pressure (Fig. S12A). The fusion to miniTurbo did not affect the plastid targeting of MpTACO1, as evident by immunoblotting from plastids isolated from the plant lines (Fig. S12B-C). We isolated biotinylated proteins and determined their identity through mass spectrometry. As expected from the dual localization, the biotinylated proteins included 56 dual-, 156 plastid- and 50 mitochondrial proteins (Fig. 5A, Table S3). On the whole, 89.1% of the biotinylated proteins were organelle localized (Fig. S12D) and were represented by higher number of unique peptides (mean=12 peptides) than non-organelle proteins (mean=7.2 peptides). On the contrary, for MpSERK (Mp7g09160) – a receptor-like kinase of the plasma membrane we used as a control^58^ – 84.6% of the biotinylated proteins were cytosolic proteins (Fig. S12D). Principal component clustered the differentially abundant proteins in the MpTACO1 and MpSERK separately (Fig. S12E), further substantiating the validity of proteins in the proximity of MpTACO1. In line with an expected role in protein synthesis, five of these proteins were organellar ribosomal subunits (e.g. bS18c, L31, uS15c, uS5c), three were involved in ribosomal biogenesis (Large ribosomal RNA subunit accumulation protein YceD), and 17 in translation (e.g. organelle translation elongation factors Tu, Ts, G, P and GUF1). Broad functional categories also underscored candidate interactors involved in photosynthesis, energetics, protein folding, degradation and membrane translocation (Fig. 5B). A comparison of the candidate interaction partners of MpTACO1 with those identified in human^39^ and mouse^40^ shows that the shared proteins are exclusively organelle localized and include ribosomal subunits (e.g. uS5m and uS15m, identified in two of the species) and elongation factors (e.g. elongation factor Ts identified in all three species). At the genome level, more than a dozen candidate interaction partners, including ribosomal subunits and elongation factors are universally conserved (Fig. 5D) and proteins involved in chloroplast protein biogenesis, import and membrane insertion are conserved across photosynthetic organisms. Together, the proteins identified by proximity labelling confirm the dual-localization of MpTACO1 and across species comparisons support its conserved role in translation in mitochondria and plastids.

**Fig. 5:**
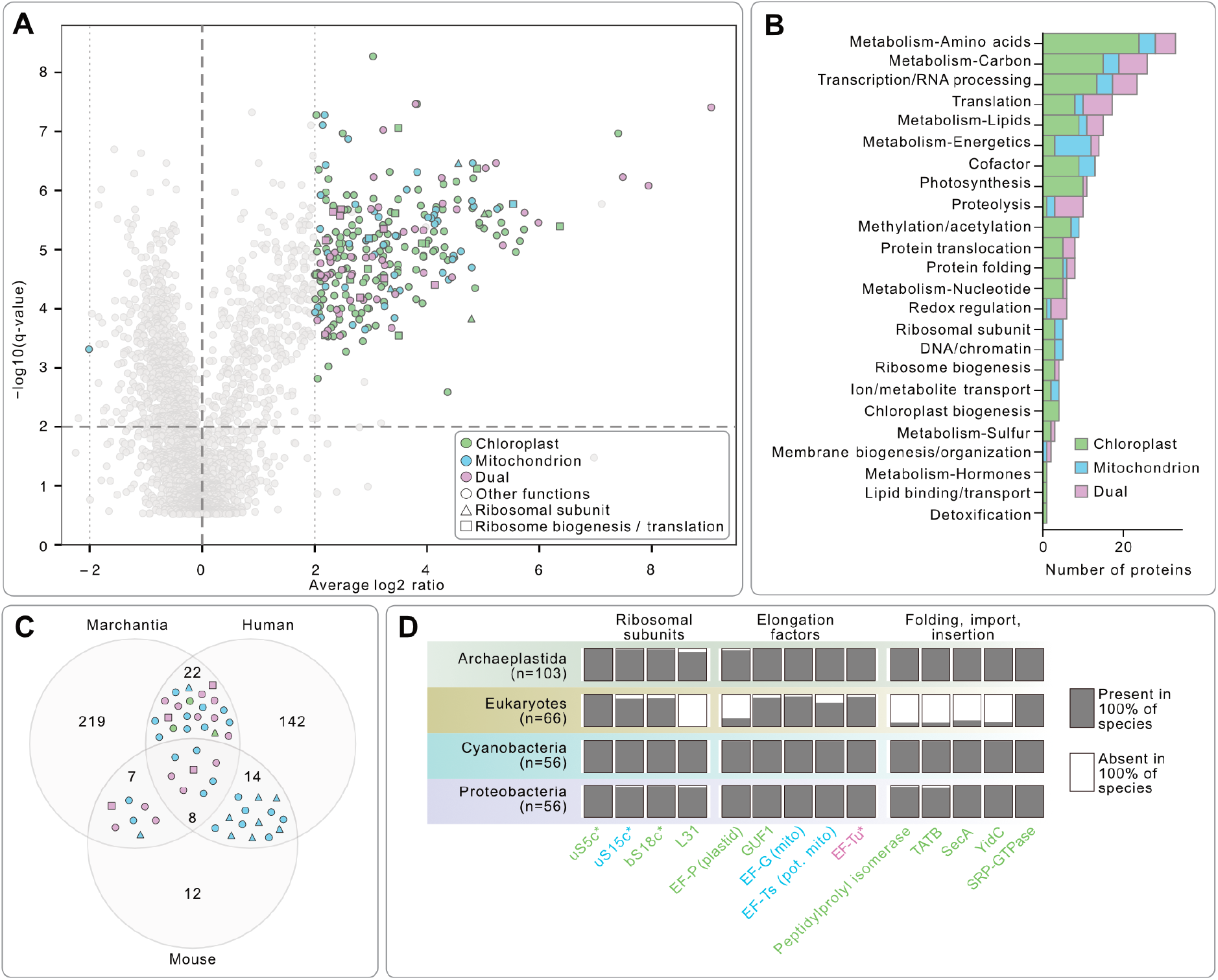
Proximity-labelling by TACO1:miniTURBO in mitochondria and plastids. **(A)** Log₂ fold change in protein abundance (MpTACO1::miniTurbo/WT) plotted against −log₁₀(q value). Colored points represent 263 organelle-localized candidate proteins; colors indicate localization and shapes indicate functional class. Light-grey points represent 2,354 non-candidate background proteins: those that did not meet the candidate criteria (q < 0.01 and |log₂FC| > 2) or were also differentially abundant in the negative control. **(B)** Functional categories of organelle-localized proteins enriched in miniTurbo lines. Stacks within each bar represent protein localization. **(C)** Comparison between experimental candidate interactomes of Marchantia (this study), mouse^40^ and human^39^; shared proteins are indicated inside the overlapping regions as per the key in A. **(D)** Evolutionary conservation of a subset of enriched proteins that are involved in information processing (transcription and translation) and photosynthesis in plastids across eukaryotes bearing plastids, other eukaryotes, cyanobacteria and proteobacteria (n, number of species in each category). Protein names are colour-coded to represent their localization as per the key in A, asterisks indicate proteins experimentally identified in TACO1 interactome of at least two species (i.e. overlapping regions in Fig. C).

To further investigate the role of MpTACO1 in the two organelles, we aimed to generate loss-of-function lines using CRISPR-Cas9^59^. We initially designed sgRNAs targeting four distinct regions coding for the targeting peptide, domain 2 and domain 3 (Fig. S13A). We transfected *Marchantia* TAK1 thalli with these sgRNAs constructs at least thrice along with Cas9 and found that on average, out of 50 explants, 3-5 regrew (Fig. S13B). They all turned out to be wild type in sequence and growth phenotype (exemplary plants shown in Fig. S13C). We then designed two additional sgRNAs targeting the 5’ UTR to introduce early stop codons and transfected *Marchantia* BoGa strain spores to increase the chances of selecting lines with successful isogenic mutations in Mp*TACO1*. This recovered two plant lines with amino acid substitutions, albeit only in the TP region. These mutant plants also resembled WT in their thalli growth, PSII photochemical efficiency, electron transport rate, quantum yield, non-photochemical energy dissipation and fraction of open reactions centres (Fig. S13D-E). Together, the failed attempts to generate loss-of-function lines in two distinct strains and developmental stages suggested that TACO1 in *Marchantia* is not readily dispensable.

## Discussion

We provide evidence for the selection and retention of a single YebC/TACO1 homolog of bacterial origin in algae and plants, which is dual targeted to mitochondria and plastids. Data from bacteria and mitochondria demonstrate the protein acts as an auxiliary protein to their ribosomes and helps to stabilize the PTC during the translation of proline motifs^37–39,41^. Organelle-encoded proteins often nucleate the early assembly of complexes for oxidative phosphorylation and photosynthesis, respectively at the inner mitochondrial or thylakoid membrane. This process requires the coordination of an import machinery, organellar translation, assembly factors, and the co-translational membrane insertion of nascent peptide chains and the addition of metal moieties such as heme or chlorophyll.

Unassembled subunits, in particular those involving metal cofactors, are problematic for mitochondria and plastids, as evident from an elaborated machinery that has evolved for their rapid degradation^60–65^ in a way that resembles the envelope stress response known from bacteria^66^. Organelle translation must hence secure the spatiotemporal translation, folding and correct insertion at their membranes. In this respect, ribosomal stalling by PP motifs could offer a regulatory mechanism. Prolonged stalling, however, is simultaneously problematic and numerous rescue factors such as EF-P, eIF5a, and the ABCF protein family emerged – likely together with the origin of cellular life itself – to deal with motifs that can slow down ribosomes^35^. These factors have been co-evolving for billions of years with their respective ribosomes in bacteria as well as eukaryotic organelles of bacterial origin. While bacterial and mitochondrial translation with respect to proline motifs and the recruitment of YebC/TACO1 proteins has been explored^37–39,41^, this is not yet the case for plastid ribosomes.

Polyproline motifs are a common cause of stalling, especially when preceded by D/A or followed by G/W/N/D ^67^. Among the more than 15,000 plastid genomes we screened, 29.4% of the encoded proteins harbour at least one PP motif. That is less than the 41.7% of cyanobacterial proteins and showcases the different evolutionary forces acting on plastid genomes that reduce the number of GC-rich codons in general^45,46^. The presence of PP motifs encoded by CCN hints at a retention for reasons beyond their ability to adopt unique right- or left-handed type I and II helices^68^. Proline residues present a dilemma for translation at the PTC due to the biochemical nature of the amino acid, and which is likely why, in fast growing bacteria, the frequency of the amino acid is reduced^35,69^. The high frequency of PP motifs retained across all domains of life, however, is testimony to their benefit or even necessity (e.g. as a helix breaker). An analysis of 43 *Escherichia coli* strains underscores the regulatory function of PP motifs^70^ that also have a tendency to increase in number in more complex eukaryotes^71^. PP motifs encoded on plastid genomes are also found in the vicinity of transmembrane domains or at C-termini (Fig. S3), which could aid (i) their insertion by insertases of the OXA family in both mitochondria and plastids^72,73^ and (ii) steps in the maturation of a protein such as the addition of metal cofactors. Thus, emerging data appears at odds with literature that view PP motifs as a mere inconvenience during translation^35^ or only as helix breakers during protein folding, as it neglects their potential to act as indicators for events in translation that require pausing^74–76^. The evolutionary retention of auxiliary proteins shows that ribosomes require them to manage the rate of translation of PP motifs encoded within the mRNA, even if ribosomes can synthesise peptides encoding PP motifs at a dramatically slower rate in the absence of additional factors under laboratory conditions^77^. This points to temporal regulation of nascent chain synthesis, folding and membrane insertion. A broad and global analysis of protein sequence features upstream of polyproline motifs could address this question and might identify different functional categories for the motifs.

Mitochondrial ribosomes require TACO1 for elongation of at least some proteins enriched in PPs, and we show TACO1 to be imported also by Marchantia plastids (Fig. 4). Proximity labelling using a miniTurbo-tagged MpTACO1 (Fig. 5) identified 56 dual targeted proteins, supporting the dual localization of MpTACO1. Enrichment of organelle ribosomal subunits, translation elongation factors, and proteins involved in protein folding (including a peptidylprolyl isomerase that accelerates protein folding at polyproline motifs), suggest an overall interactome profile similar to that previously identified in mitochondria and a similar function of MpTACO1 in plastids. The enrichment of highly conserved proteins involved in protein import and membrane insertion (e.g. protein translocase TATB, secA and YidC; chloroplastic Signal-recognition-particle GTPase and TIC 40) places MpTACO1 in the vicinity of the inner plastid membrane and the thylakoids. While the interaction of each of these candidates, and their functional relevance, remains to be directly tested, the overall profile support dual targeting as well as a role in protein synthesis.

Like Marchantia, and the vast majority of plants, maize and Arabidopsis encode a single TACO1 homolog (S1B) and public databases such as TAIR list the protein as plastid-localised. The single copy conservation of a YebC/TACO1 contrasts with the otherwise observed expansion of organelle-related protein families in land plants^47,78^. Encoding YebC/TACO1 paralogs seems unfavourable, both in pro- and eukaryotes, and one YebC copy was selected against early in the evolution of the first photosynthetic eukaryote – the primal alga – before the diversification into the three main archaeplastidal lineages (Fig. 1A, Fig. S1B). Phylogenomics was not able to resolve which one (Fig. 1C), but we speculate the cyanobacterial YebC, since the mitochondrial TACO1 was already nuclear-encoded with a functional targeting sequence. Dual targeting to mitochondria and plastids is observed more frequently for proteins of transcription and translation^55^ and thought to require a combination of balanced charges, serine residues and a mitochondrial membrane potential^54,79,80^. Research on dual-targeting outside of Arabidopsis is meagre, reporter-based studies are rare, and algorithm-based predictions of dual targeting error-prone^81^. Our growth conditions and detection limits did not capture a localisation shift for the serine-to-alanine mutants, adding further mystery to the role of the significant enrichment of serine residues in pTPs. The three transcription and translation-associated proteins that we show are dually targeted in the bryophyte in addition to TACO1 (Fig. S11), however, support the described pattern for Arabidopsis^55^ and can help to experimentally explore the extent, evolution, and regulation of dual targeting to mitochondria and plastids.

The structure of the first plant TACO1 homolog, in combination with a bioinformatic analysis, underscores a level of conservation that matches that of ribosomal proteins. It speaks for the concatenated evolution of the accessory protein that is recruited to the peptidyl transfer centre (PTC) when both A- and P-sites are occupied by prolyl-tRNAs^38^, ever since its origin in the last common ancestor of bacteria^35^. Accordingly, our docking simulation results show that MpTACO1 could have a similar interaction interface with Marchantia chlororibosome bL12 to that found in the human mitoribosome-TACO1 structure. This high structural conservation points to a similar function of MpTACO1 in the chlororibosome (Fig. 3). The closed conformation from the SAXS data is consistent with the cryoEM reconstruction of the HsTACO1 that is bound to mitoribosomes and the flexibility we demonstrated is consistent with the molecular dynamics results^43,44^. However, additional structural data would be needed to conclusively show that this is the case.

Thus, localisation-, structural-, proximity labelling-, and phylogenomic data point to the plant YebC/TACO1 having a similar function in the plastids of algae and plants as its homolog in the mitochondria of non-photosynthetic eukaryotes^40,43^ and the cytosol of bacteria^38,39^. This extends the range of substrates, whose translation is enhanced by eukaryotic YebC/TACO1 homologs, supporting its importance for the translation of bioenergetic membrane proteins of endosymbiotic organelles. MpTACO1’s crucial role in the translation of organelle-encoded proteins likely explains why it is not easily mutated, except for amino acid substitutions in the targeting peptide. This suggests that, while CRISPR-Cas9 is functional, cells with mutations rendering the ribosome’s accessory protein non-functional (e.g. early stop codons) are selected against when plants are regrown from gemmae or sporeling stages —i.e. early developmental stages, where MpTACO1 is relatively highly expressed (Marchantia.info). Loss of TACO1 in human and mouse also results in severe phenotypes^40–42^ and its role in both organelles likely exacerbated such phenotypes in Marchantia. The current data hence suggests that this single copy, highly-conserved gene, is likely essential for plants.

## Conclusion

The bioenergetic membranes of mitochondria and plastids are of bacterial origin. The maturation of their protein complexes requires co-ordination of translation, modification and insertion of a variety of proteins such as COX1 into the inner mitochondrial membrane or D1 (psbA) into the thylakoids of plastids. Both these proteins contain conserved PP motifs that are also frequent across other organelle-encoded proteins, and many of which are evolutionary conserved and known to recruit accessory proteins to the ribosome during translation. YebC/TACO1 is one such protein required by mitochondria and its targeting to plastids suggest a similar role in their translation of at least a subset of about two dozen PP-containing proteins underscored here. As for the presence of PPs, we speculate that the retention of PP motifs encoded by GC-rich codons in AT-rich genomes also hint at a regulatory potential in translation elongation for nascent chain folding and membrane insertion. While the mechanism of the YebC/TACO1 recruitment to ribosomes remains uncertain, one can synthesize a model covering bacteria, mitochondria and plastids. The PTC site of their ribosomes is constantly probed by YebC/TACO1 proteins that are rapidly changing between conformations. Formation of peptide bonds involving prolines is slower, triggering a conformational shift in domain V of the 23S rRNA (or its mitochondrial equivalent, the 16S rRNA), which recruits YebC/TACO1 that can then interlock in its closed conformation to stabilize the PTC site. We propose the protein family name be changed to <u>T</u>ranslation <u>E</u>nhancer of (<u>P</u>oly)<u>P</u>rolines (TEPP) proteins based upon the following data: (a) the localisation of TACO1 to plastids suggest a role in translation regulation beyond the mitochondria-encoded COX1; (b) the factors bind primarily rRNA, not mRNA, across different ribosomes; (c) the factors regulate the rate of translation elongation and not activate translation initiation; and (d) the robust structural and functional conservation since YebC origin. Our data hint at a so far untapped feature of plastid biology, which is proline residue translation that requires ancient accessory ribosomal proteins of the TEPP family such as YebC or TACO1.

## Methods

### Comparative genomics

We searched for poly-proline regions (PPs; defined as at least two consecutive prolines in protein sequence) across all chloroplast and mitochondrial genomes available on NCBI-Refseq (accessed in March 2026) using an inhouse python script (‘count_PPs.ipyub’). To search for ribosomal rescue factors across species, we clustered whole proteome models of 169 eukaryotic, 57 proteobacterial and 56 cyanobacterial genomes using OrthoFinder version 2.5.4^82^ into protein orthologues (Table S4) and identified ribosomal rescue factors using *Arabidopsis*, Human or *E. coli* gene IDs of rescue factors reported previously^35^. For generating a phylogenetic tree of TACO1/yebC homologs, we aligned protein sequences of 13 diverse eukaryotes, 7 diverse cyanobacteria and 8 diverse proteobacteria using MAFTT (v7.525)^83^, and removed the N terminal targeting peptide extension (region that did not align with prokaryotic homologs) from the eukaryotic homologs before generating a phylogenetic tree and predicting structures. The phylogenetic tree was generated using IQTree v1.6.12^84^ and structures were predicted using AlphaFold3^48^. The models with highest confidence were used for structural comparison on DALI^85^.

### MpTACO1 expression, purification and crystallisation

A recombinant plasmid containing the MpTACO1 coding sequence (Mp1g23800) was constructed using Gibson assembly. The coding sequence excluding the predicted mitochondrial targeting sequence was amplified from pDONR221_MpTACO1_NoNTS using Mar_For (ACAGAGAACAGATTGGTGGAATGGGGAGACGTTCGGCCAAG) and Mar_Rev primers (ACTCGAGTGCGGCCGCAAGCTTATTACTTTTGATTACAATAGACAGCATCCACA).

Polymerase chain reaction (PCR) amplification was performed with the Platinum SuperFi II Green PCR Master Mix (Invitrogen) with 0.5 µM primers and 2 ng of template DNA. The backbone plasmid for the Gibson assembly was pFRGSF01^86^, which encodes for both a hexa-histidine tag and a SUMO tag at the N-terminus. pFRGSF01 was amplified using primers pFR_For (GCTTGCGGCCGCACTCGAGT) and pFR_Rev (TCCACCAATCTGTTCTCTGTGAGTCTCA) with the Platinum SuperFi II Green PCR Master Mix (Invitrogen), 0.5 µM primers, and 10 ng of template DNA. The PCR product was digested with DpnI (Thermo Scientific). Both PCR products were purified using a NucleoSpin Gel and PCR Clean-up kit (Macherey-Nagel). For the Gibson assembly, a NEBuilder HiFi DNA Assembly Cloning Kit (New England BioLabs) was used. The assembly reactions were transformed into XL10 Gold electrocompetent (E. coli cells and incubated overnight at 37 °C on Luria-Bertani agar supplemented with 25 µg mL-1 kanamycin.

For expression, the recombinant plasmid was transformed into E. coli BL21(DE3) and the gene was expressed in autoinduction media (AIM-LB broth base including trace elements, Formedium). Cultures were grown at 37°C with shaking until they reached an optical density (OD600) of 0.6, after which the temperature was reduced to 20°C for 16h. Cells were collected by centrifugation (2719 RCF Rmax, 10 min, 4°C). The cell pellets were resuspended in 50 mL 20 mM Tris-HCl pH 8.0, 150 mM NaCl, 30 mM imidazole pH 8.0 supplemented with 1 tablet of Pierce™ Protease Inhibitor Tablets, EDTA-Free (Thermo Scientific) and then lysed using an EmulsiFlex-C3 (Avestin). Cell debris was removed by centrifugation (26712 RCF Rmax, 30 min, 4°C). The clarified lysate was incubated with immobilized metal ion affinity chromatography Sepharose™ 6 Fast Flow resin (GE Healthcare) pre-charged with 0.1 M NiSO_4_. The resin was washed with five column volumes of 20 mM Tris-HCl pH 8.0, 150 mM NaCl, and 30 mM imidazole pH 8.0, then packed into a disposable PD-10 gravity column (Cytiva) and washed again with five column volumes of the same buffer. Bound proteins were step-eluted with 20 mM Tris-HCl pH 8.0, 150 mM NaCl, 200 mM imidazole pH 8.0, and collected as 1 mL fractions. The fractions with the highest protein concentration and reasonable purity were combined and diluted with 20 mM Tris-HCl pH 8.0, 50 mM NaCl cleaved with SUMO protease on ice for 1 hour. Further purification on a HisTrap™ FF crude 5 mL column and HiTrap™ Q HP 1 mL (Cytiva) column in tandem, prior to elution from the The HiTrap™ Q column with a linear gradient from 50 mM Tris-HCl pH 8.0 to 100% 50 mM Tris-HCl pH 8.0, 1M NaCl. Finally, the peak fractions were polished on a Superdex™ 75 10/300 GL gel filtration column (Cytiva) equilibrated with 50 mM Tris-HCl pH 8.0, 150 mM NaCl.

Crystallisation experiments were conducted in the FINStruct HiLIFE Crystallography Unit at the Institute of Biotechnology, University of Helsinki, using a mosquito® LCP (SPT Labtech). Purified recombinant MpTACO1 was concentrated to 13.7 mg mL^-1^, using Sartorius Vivaspin® 500 Centrifugal Concentrator (10,000 MWCO). Crystallisation was carried out by sitting drop vapour diffusion at 20°C, with 1:1 (v/v) mixtures of protein sample and crystallization solution (100 nL each). The crystallisation plates were imaged using a Rock Imager 182 (Formulatrix). Diffracting plates were obtained after 20 h from the Helsinki cryoscreen (https://www.helsinki.fi/en/infrastructures/integrated-structural-cell-biology/infrastructures/crystallization/standard-screens, accessed 17.3.2025) from a main solution of 0.1M MES, pH 6, 30% (v/v) polyethylene glycol 200, 5% (w/v) polyethylene glycol 3350. Data was collected at beamline ID30A-3 (ESRF, Grenoble, France) at a wavelength of 0.9677 Å. The X-ray diffraction data were auto-processed with the GrenADES parallelproc pipeline. The Alphafold3^48^ model without the predicted MTS was used for molecular replacement using Phaser^87^ as implemented in the CCP4 Cloud ^88^. The structure was initially refined using *REFMAC5*^89^ and ModelCraft^90^, followed by several rounds of manual building in *Coot*^91^ with further refinement and validation in PHENIX^92^. Figures were prepared in UCSF ChimeraX^93^

### Small angle X ray scattering (SAXS) analyses on MpTACO1

We collected the SAXS data on our Xeuss 2.0 Q-Xoom system from Xenocs, equipped with a PILATUS 3 R 300K detector (Dectris) and a GENIX 3D CU Ultra Low Divergence x-ray beam delivery system. The chosen sample to detector distance for the experiment was 0.55 m, results in an achievable q-range of 0.05 – 5.5 nm^-1^. The measurement was performed at 15°C with a protein concentration of 5.44 mg/ml. The sample was injected in the Low Noise Flow Cell (Xenocs) via autosampler. We collect 36 frames for buffer and protein with an exposer time of ten minutes/frame and scaled the data to absolute intensity against water. Data frames were checked for radiation damage and averaged, leading to buffer subtracted protein data. All used programs for data processing were part of the ATSAS Software package (Version 3.0.5)^94^. Primary data reduction was performed with the program PRIMUS^95^. With the Guinier approximation^96^, we determine the forward scattering *I(0)* and the radius of gyration (*R_g_*). The program GNOM^97^ was used to estimate the maximum particle dimension (*D_max_*) with the pair-distribution function *p(r)*. We compared the solved crystal structure with the experimental data using CRYSOL^98^. A Model was created with AlphaFold3^48^ to complete the crystal structure. The helical N-terminal part of the model (domain 1) was set as flexible and remodeled with an Ensemble Optimization Method (EOM)^99,100^. The resulting EOM models were checked for sterical clashes, and the YASARA Energy Minimization Server was used to improve these models^101^.

### Modelling and docking simulations

The Consurf web server was used to plot conservation^51,52^. PDB files were prepared using the PDB-Tools web server (https://wenmr.science.uu.nl/pdbtools/) before docking in HADDOCK3^53^. Potential active residues involved in interactions at the interface between bL12 and TACO1 were selected based on the human structure (PDB ID 9OLF^43^, Fig. 3 c-d). Homologous active residues in *M. polymorpha* for both proteins were identified and HADDOCK3^53^ was used to identify potential passive interacting residues in proximity of the interface. Active and passive residues were included to make ambiguous restraints, with all passive residues of one protein being specified as potential interaction partners for active residues of the other protein.

### Plant transfectant generation and growth conditions

All transfectants in this study were generated in ecotype Takaragaire-1 (Tak-1) of the liverwort *Marchantia polymorpha*. The wild type gemmae were grown under continuous light (70 μmol m^-2^ s^-1^) on half-strength Gamborg B5 vitamin agar-plates (GVA). We amplified the genes of interest from *Marchantia,* Mp*TACO1* (Mp1g23800 alias Mapoly0061s0140), aspartyl/glutamyl amidotransferase subunit B (Mp1g29210 alias Mapoly0107s0036), lysyl-tRNA-synthetase (Mp7g09810 alias Mapoly0003s0001), seryl-tRNA-synthetase (Mp6g09660 alias Mapoly0016s0010), LHCA3/LHCII Chla/b binding protein 3 (Mp7g08940 alias Mapoly0068s0047) and MME1 (Mp4g02270 alias Mapoly0080s0072) from the cDNA of 2 week old thalli and cloned into *Gateway* (Invitrogen) entry vector pDONR221^TM^ (Invitrogen) by BP reactions set as per manufacturer’s protocol. Substitutions of interest were made into the wild type Mp*TACO1* using PCR-mutagenesis. All constructs were transferred N-terminal of Citrine into pMpGWB106 ^102^ via *Gateway* cloning by LR reactions set as per manufacturer’s protocol.

We integrated the localization vectors with the genes of interest into Marchantia genome via *Agrobacterium* mediated transfections previously optimized^54,103^. Briefly, the vectors were transfected to electrocompetent *Agrobacterium tumefaciens* (GV3101 lacking pSOUP) by electroporation (Bio-Rad GenePulser Xcell, 1.44 kV) and *Agrobacterium* transfectants were incubated with explants from two-week-old thalli after removing their epical notches and two days of recovery. Transfectants were then selected onto Hygromycin (10 μg/mL) and Cefotaxime (100 μg/mL) and further select for successful reporter expression at gemmae under the microscope. Expression positive gemmae were further propagated and used for the localization experiments after growing for up to three generations by serial passages from the gemma stage.

### Sample preparation for subcellular localization

Localization experiments were conducted in gemmae, 2-day old thalli or protoplasts isolated from 4-7 days (until otherwise stated) old thalli of isogenic transfectant lines. Gemmae or thalli were placed on a glass slide with 30-50µL water and covered with a cover-slip. Protoplasts were isolated as previously described^54^, from the thalli after incubation with 10mL of 8% Mannitol, 6.3 g/L Gamborg B5 Vitamins (MGV) with 15mg/ml Cellulase and 5 mg/mL Macroenzyme R10 and filtration with a 50µm strainer. After two washes with MGV (at 300g, 3min), the protoplasts were used for mitochondrial staining by MitoTracker™ Red CMXRos (following the manufacturer’s protocol).

### Microscopy

Microscopy was carried out as described previously^54^. Briefly, protoplasts were imaged using Nikon eclipse Ti imaging platform (illuminated with Nikon Intensilight C-HGFIE for fluorescence, connected to the camera via Nikon Digital Sight DS-U3 and to the PC via NIS-Elements Basic Research version 4.30) for visualizing chloroplast autofluorescence (480/590 nm, exposure time 20-80ms), Citrine (470/520 nm, exposure time up to 300ms) and mitochondria (540/605, exposure time up to 300ms). To control for plastid autofluorescence during Citrine readout, the mitochondrial malic enzyme (MME1) was fused with citrine, expressed and imaged under identical conditions (images in Fig. S11). 2-day old thalli (in Fig. 3A and 3C) were imaged on a Leica TCS SP8 platform (Leica, Wetzlar, Germany) with an HC PL APO CS2 93× glycerol objective (NA 1.3, Leica), connected to a PC via LAS-X (Leica) and deconvoluted using the Lightning adaptive deconvolution module. Plastid autofluorescence was filtered out while scanning for the Citrine signal using time gating^56^. The collages were prepared in ImageJ (Fiji). Co-localisation analyses were conducted using the BIOP JACoP plugin.

### Plastid isolation

Plastids were isolated following the lysis of 2-week-old thalli, in presence of 5-10 mL isolation buffer (50mM HEPES KOH pH 7.5, 0.33 M sorbitol, 1mM MgCl_2_, 1mM MnCl_2_, 2mM EDTA pH 8, 0.5% w/v BSA, 0.5% w/v PVP40, and freshly added 5mM ascorbate), the lysate was filtered through a Mira cloth and centrifuged at 1000g, 10min. The pellet was resuspended in IB and loaded onto a discontinuous gradient of 70-30% Percoll. Followed by density gradient centrifugation at 4300g, intact plastids were harvested from the 70-30 layer, washed with wash buffer (WB; IB without BSA and PVP40) three times by centrifugation for 10min at 3000g, 200og and 1000g. Plastids were pulverized in presence of liquid nitrogen and crude plastid lysate was used for immunoblotting.

### Interaction partners of MpTACO1

We investigated interaction partners of MpTACO1 using mini-turbo based method optimized previously for *Marchantia*^57^. Briefly, we fused the full length MpTACO1 at the N terminal of miniTurbo protein onto vector pMKMM2 via *Gateway* cloning by LR reactions set as per manufacturer’s protocol. The final constructs were transfected to Marchantia as per above and transfectants were selected on 0.5µM Chlorsulfuron for two generations and used for the biotinylation assay in the third generation. For biotinylation, gemmae (10-12 gemmae per plate, four plates per replicate) were grown for 10 days, and thalli were soaked into 600µM biotin (ca. 2-3 thalli per 10mL biotin solution in water) subjected to 5 min of vacuum infiltration and incubated further for 18 hours. All thalli from a single replicate were pooled and washed three times with 50ml water and tapped briefly on tissue paper for the removal of water prior to lysis. Dried thalli were pulverized in liquid nitrogen, the powder resuspended in SDT buffer (100 mM Tris, pH 7.5, 4% SDS, and 0.1 M DTT), incubated at 45 °C for 15 min for resuspension. Lysate was cleared by two rounds of centrifugation at 10,000 g 15 min, and used for protein isolation. To isolate protein, 666µL methanol was mixed well by vortexing with 500 µL lysate, mixed further with 166µL chloroform and 300µL water. After centrifuge at 10,000 g 10 min the upper and lower liquid layers were removed to retain only the middle proteinaceous middle layer in the tube. This layer was washed with 600 µL methanol by mixing well with pipetting and sonication in ultrasonic bath for 10 min. This was centrifuged at 10,000 g, 10 min and washed again. Methanol was removed by decanting after the second wash and by leaving the tube upside down for 1-2minutes. The pellet was resuspended in 500µL SDT at 37°C for 10min while shaking at 1000rpm, sonicated in ultrasonic bath for 10min, and again incubation at 37°C for 10min while shaking at 1000rpm. The resuspended proteins were used for affinity pulldown.

100µL of 50% Streptavidin agarose bead slurry was mixed with 5mL binding buffer (0.1M phosphate, 0.15M NaCl, 0.5% SDS, pH 7.2), mixed by inverting 5-10 times and centrifuged for 2min at 3500g at room temperature (this step was repeated three times). 500µL of isolated protein was added to 3.5mL of wash buffer 2 (0.1M phosphate, 0.15M NaCl, pH7.2) and transferred to streptavidin beads and mixed by inverting 5-10 times. After 18h incubation at 22°C (gentle rocking) beads were centrifuged at 3500g 3min and removal of as much supernatant as possible. The beads were washed four times with wash buffer 1 (0.1M phosphate,0.15M NaCl, 2% SDS, pH 7.2). The beads were incubated in Laemmli buffer, centrifuged and supernatant was resolved on SDS-PAGE and individual gel pieces were reduced with 50 µl 10 mM DTT, alkylated with 50 µl 50 mM iodoacetamide and digested with 200ng trypsin in 100 mM ammonium bicarbonate. Peptides were subjected to liquid chromatography after resolving them in 15 µl 0.1% trifluoroacetic acid. LC-MS analysis was performed using an Evosep One liquid chromatography system (Evosep, Odense, Denmark) coupled online to a timsTOF Pro 2 mass spectrometer (Bruker Daltonics, Bremen, Germany). Peptide separation was performed using a 15 cm Aurora Elite column (IonOpticks, Australia) in Whisper Zoom configuration at a throughput of 20 samples per day.

Raw files were processed using Spectronaut version 19.8 (Biognosys) against Marchantia polymorpha whole nuclear, mitochondrial and plastidal genome models (FASTA file provided with the figure source data). Searches used trypsin specificity, with carbamidomethylation (C) as a fixed modification and oxidation (M) and protein N-terminal acetylation as variable modifications. Mass tolerances and other search parameters were set to the software default settings. Precursor and protein identification thresholds were set to a Q-value cutoff of 0.01. Q-values were calculated using the target-decoy approach implemented in Spectronaut. Protein abundances were quantified based on MS2 fragment ion peak areas using the automatic protein LFQ method. Cross-run normalization was enabled, with normalization strategy and row selection set to automatic. Missing quantitative values were imputed using background signal.

The experiment comprised four biological replicates per strain (approximately 20 thalli grown from gemmae per replicate). Protein abundance differences were calculated and exported by Spectronaut (unpaired Welch’s t-test; Q-values corrected across the experiment). For each bait/WT comparison, proteins with Q < 0.01 and |log₂FC| > 2 were considered differentially abundant for each bait; proteins differentially abundant in both baits were removed to get MpTACO1-specific candidates. Marchantia proximity-labelling candidates were compared with previously identified proximity-labelling candidates of human^39^ (Supplementary Tables S2 and S3 of the original study) and mouse^40^ TACO1 (Supplementary Fig.3b of the original study). Their KEGG gene IDs are compiled in the Source Data for Figure 5C. Orthologue clusters across species (Table S4, see method section ‘Comparative genomics’) were used to find shared proximity-labelling candidates; i.e. if the same orthogroup contained proteins identified as both Marchantia and human, that protein family was considered shared and listed in the Source Data Figure 5C with the protein name or description in both species. Intracellular localisation of Marchantia candidates was retrieved from previously reported Marchantia chloroplast proteome and predicted mitochondrial proteome^81,104^. Localisation of human and mouse proximity-labelling candidates was retrieved from Mitocarta3.0^105^. The genomic presence-absence of each of the Marchantia proximity-labelling candidates was also investigated similarly and across species distributions of select proximity-labelling candidates were plotted in Fig. 5D.

### Loss of function lines for MpTACO1

The sgRNA target regions across *MpTACO1* coding sequence and 5’UTR were selected using CasFinder tool on Marchantia database (https://marchantia.info/tools/casfinder/), were cloned into pMpGE_En03 vector and subsequently into pMpGE010. Agrobacterium mediated transfection of *Marchantia* strain TAK1 thalli and *Marchantia* strain BoGa spores was conducted as described previously^103,106–108^. Briefly, epical notches of twelve-day old thalli were cut and the explants were regrown for two days before co-cultivating them for three days with Agrobacterium. Explants were washed and plated on selection medium (with Hygromycin 10mg/L). Gemmae from explants that regrew, were plated and maintained as independent transfectant lines for up to four generations. *MpTACO1* regions surrounding sgRNA target sites were sequenced in the third or fourth generations and if no mutation was observed after four generations, the experiment was terminated. For transfection of BoGa spores, each spore capsule was resuspended into 150µL of sterilization solution (0.2% sodium hypochlorite, 0.1% Triton-X100), washed with sterile water thrice, and added to spore media (1/2 Gamborg, 10% cas amino acids, 10% glutamine) and grown for seven days before co-cultivating with Agrobacteria for 3 days. Sporelings were washed and plated on selection medium similar to thalli transfection, and regrowing plant lines were maintained and screened for up to four generations. The photosynthetic parameters of PSII were measured on an IMAGING-PAM (M-Series; Walz), as described previously^107^.

## Data availability

Supplementary figures and tables 1-2 are available with this submission. Additional data are available on Zenedo (10.5281/zenodo.24222693), which includes, but is not limited to, the supplementary data, source data for the main and supplementary figures, supplementary tables, and in-house scripts. The X-ray crystallographic data are deposited with the wwPDB ID: 9SCZ and the SAXS data to the Small Angle Scattering Biological Data Bank (SASBDB)^109^ with the accession code SASDYA7. Raw data for Mass spectrometry from the proximity labelling experiment is deposited on PRIDE (Project accession: PXD087378; DOI: 10.6019/PXD087378).

## Author contributions

PKR: Conceptualization, Experimental design, Methodology, Investigation, Data curation, Formal analysis, Validation and Visualization, Writing - original draft, review and editing. NKT: Experimental design, Methodology, Investigation, Data curation, Formal analysis, Validation and Visualization CM: Experimental design, Methodology, Investigation, Data curation, Formal analysis, Visualization, Writing - review and editing MLQ: Investigation; Data curation, Visualization, Writing - review and editing TK: Investigation; Data curation DWD: Investigation, Data curation JR: Methodology, Investigation, Data curation, Formal analysis, Validation and Visualization, Writing - original draft, review and editing SHJS: Supervision; Experimental design; Funding and Resource acquisition SJB: Supervision; Experimental design; Funding and Resource acquisition, Writing - review and editing BJB: Supervision; Experimental design; Funding and Resource acquisition; Writing - review and editing. SBG: Conceptualization; Project administration; Supervision; Experimental design; Funding and Resource acquisition; Data visualization; Writing - review and editing.

## Funding

SBG acknowledges funding from the DFG (581702633) and the HHU through its strategic research fund. SJB is grateful for support from the University of Helsinki Boost funding and the Sigrid Jusélius Foundation (260021). BJB received funding from the Sigrid Jusélius Foundation Senior Investigator Award, Research Council of Finland (357469), the Magnus Ehrnrooth Foundation, and the University of Helsinki Boost funding. CM is a fellow of the MACS graduate programme, University of Helsinki. The Center for Structural Studies is part of StrukturaLINK Rhein-Ruhr which is funded by the Deutsche Forschungsgemeinschaft (DFG Grant number 573727698) and INST 208/761-1 FUGG.

## Acknowledgements

PKR and MLQ are grateful to Prof. William F. Martin for providing financial support. We acknowledge support from the high-performance computing cluster (HILBERT) HHU-Düsseldorf and the Molecular Proteomics Laboratory at HHU for conducting the mass-spectrometry. We also thank Sabine Zachgo and Nora Gutsche from the University of Osnabrück (Germany) for providing us with spores and Alice Ochs for spore transfection experiments. The facilities and expertise of the HiLIFE Crystallization unit at the University of Helsinki, a member of FINStruct and Biocenter Finland are gratefully acknowledged, as are the European Synchrotron Radiation Facility for provision of synchrotron radiation facilities under proposal ID MX2640 and on beamline ID30A-3. Molecular graphics and analyses were performed with UCSF ChimeraX, developed by the Resource for Biocomputing, Visualization, and Informatics at the University of California, San Francisco, with support from National Institutes of Health R01-GM129325 and the Office of Cyber Infrastructure and Computational Biology, National Institute of Allergy and Infectious Diseases.

## Notes

https://zenodo.org/records/24222693

### Competing Interest Statement

The authors have declared no competing interest.

### Summary of Updates

Figures 3 and 5 revised as per peer review round one; results, discussion and methods revised accordingly

