## Supplementary Figures 1-13 for "Structure determination and dual targeting of a plant TACO1 identifies its ancient role as an organelle translation regulator"

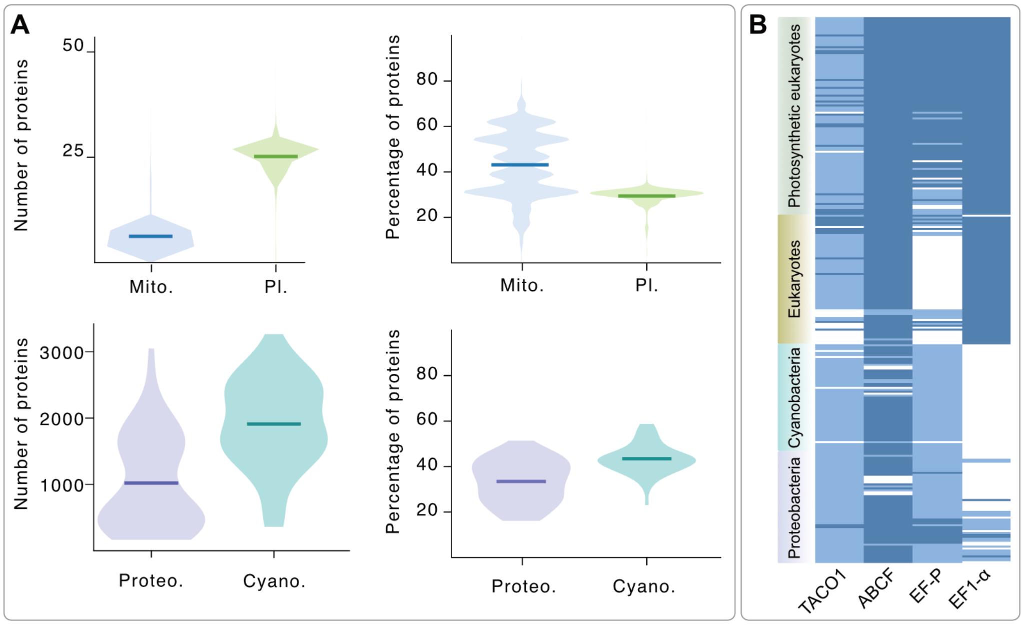

**Fig. S1: Prevalence of polyproline motifs and ribosomal rescue factors across species. (A)** Number and percentage of proteins with at least one polyproline (PP) motif across 18,059 mitochondrial, 15,208 plastid, 57 proteobacterial, and 56 cyanobacterial genomes. **(B)** Distribution of ribosomal rescue factors across 169 eukaryotic, 57 proteobacterial and 56 cyanobacterial genomes. White, no homologue; blue, single homolog; dark blue, multiple homologs.

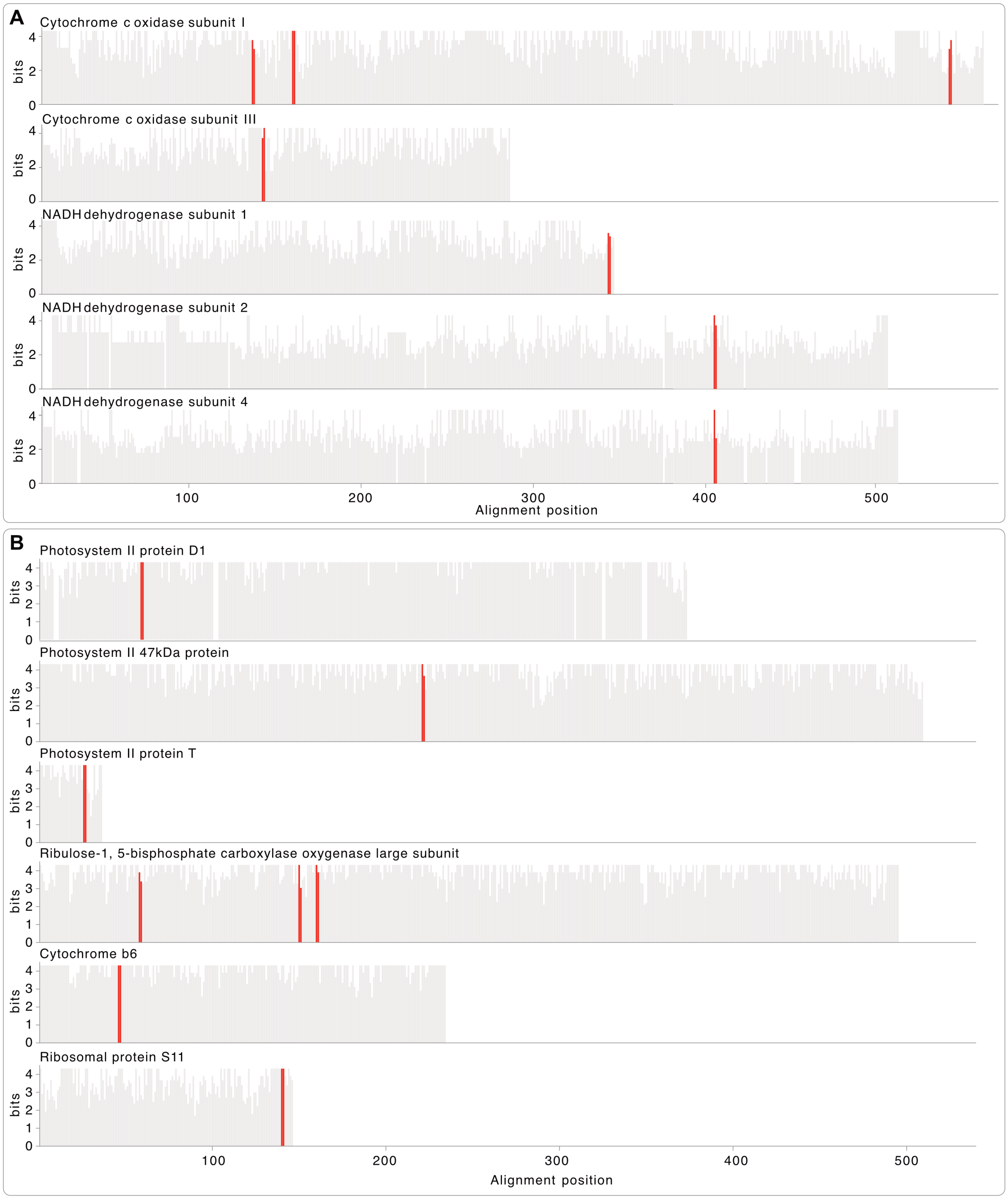

**Fig. S2: Conservation of polyproline motifs across species. (A)** Amino acid conservation across key proteins from seven mitochondrial and *Rickettsia prowazekii* (a phylogenetic relative of the mitochondrial ancestor), and **(B)** across ten plastid and *Synechococcus elongatus* (a representative cyanobacterial genome) genomes. Each bar shows the conservation of a single residue in the alignment as bit score (4 or higher = single amino acid is conserved across all species; 2, moderate conservation; 0, near-random amino acid is tolerated in that position). Red bars represent two subsequent prolines.

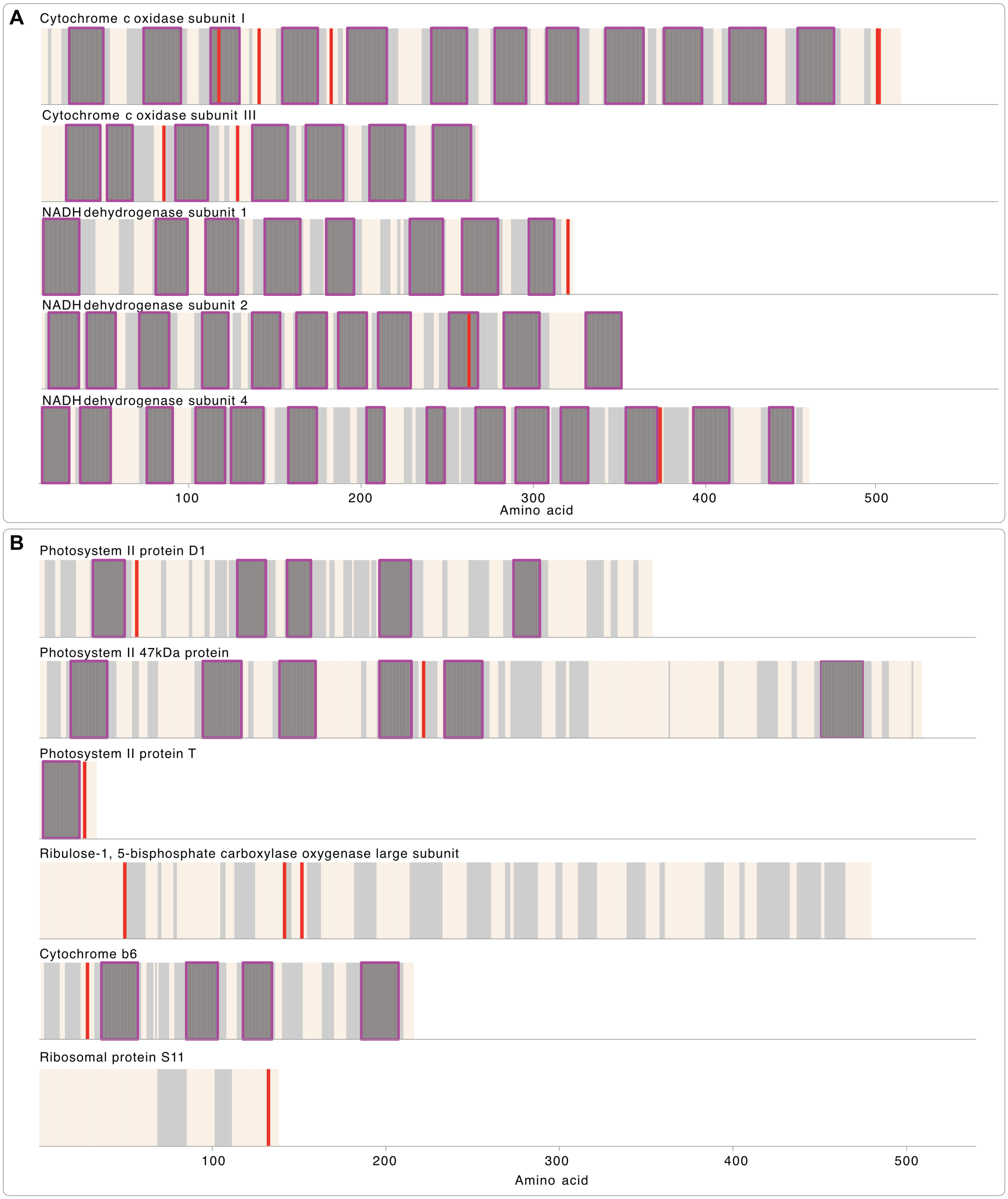

**Fig. S3: Polyproline motifs in the context of protein secondary structure of organelle encoded proteins. (A)** Secondary structure prediction for key genes from human mitochondrial and **(B)** Arabidopsis plastid genomes. Amino acids that are predicted to be inside an alpha helix (NetSurf3) are shown in light grey, those inside a membrane helix (DeepTMHMM) are in dark grey (highlighted by violet boxes), poly-prolines in red, and other amino acids in off-white.

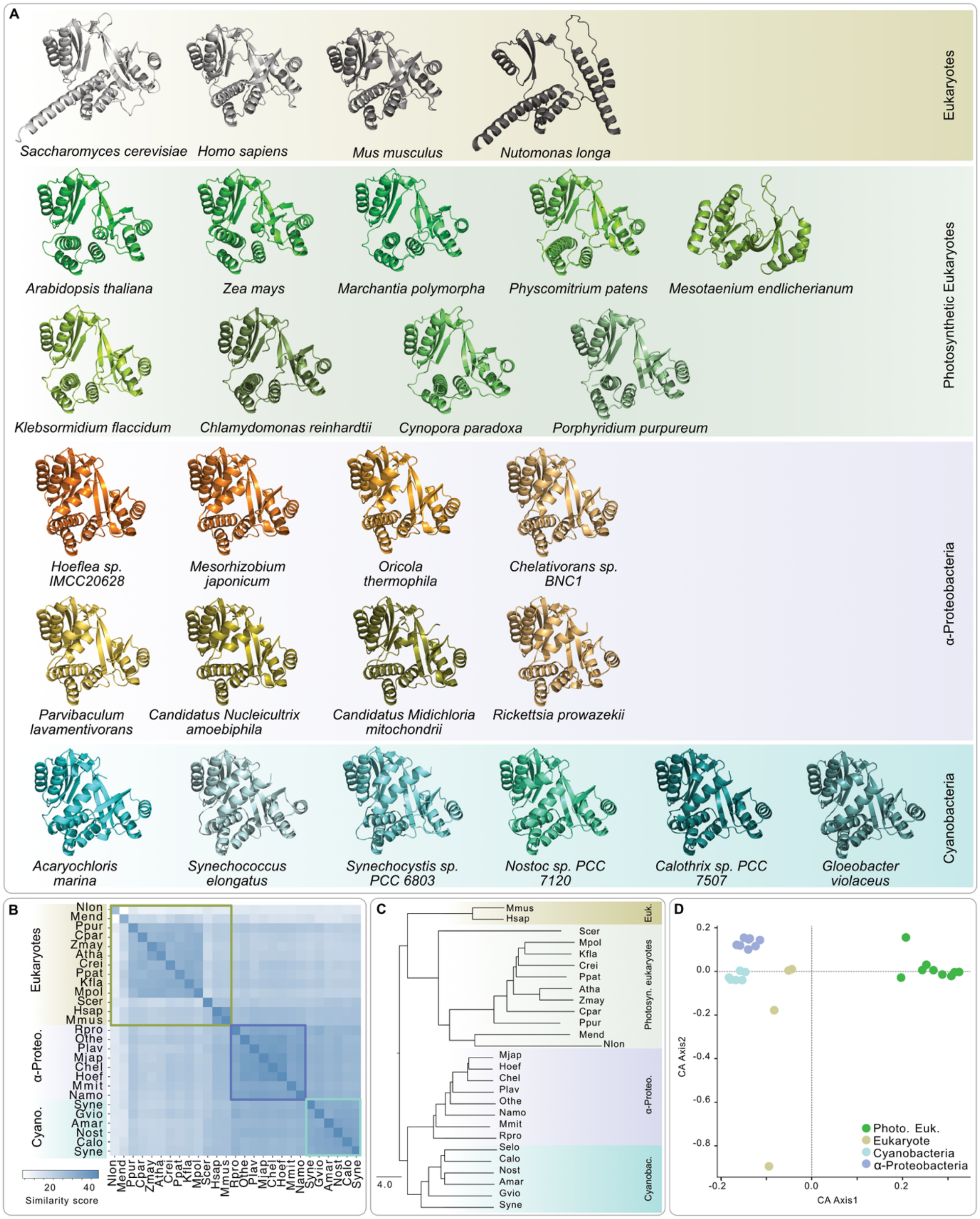

**Fig. S4: Structural conservation of TACO1 homologs across species. (A)** Predicted structures (AlphaFold 3) of TACO1/yebC homologues from eukaryotes and prokaryotes, **(B)** their DALI (Holm et al. 2020) similarity matrix, **(C)** structural similarity dendrogram rooted between prokaryotes and eukaryotes, **(D)** and a correspondence analysis that clusters similar structures together.

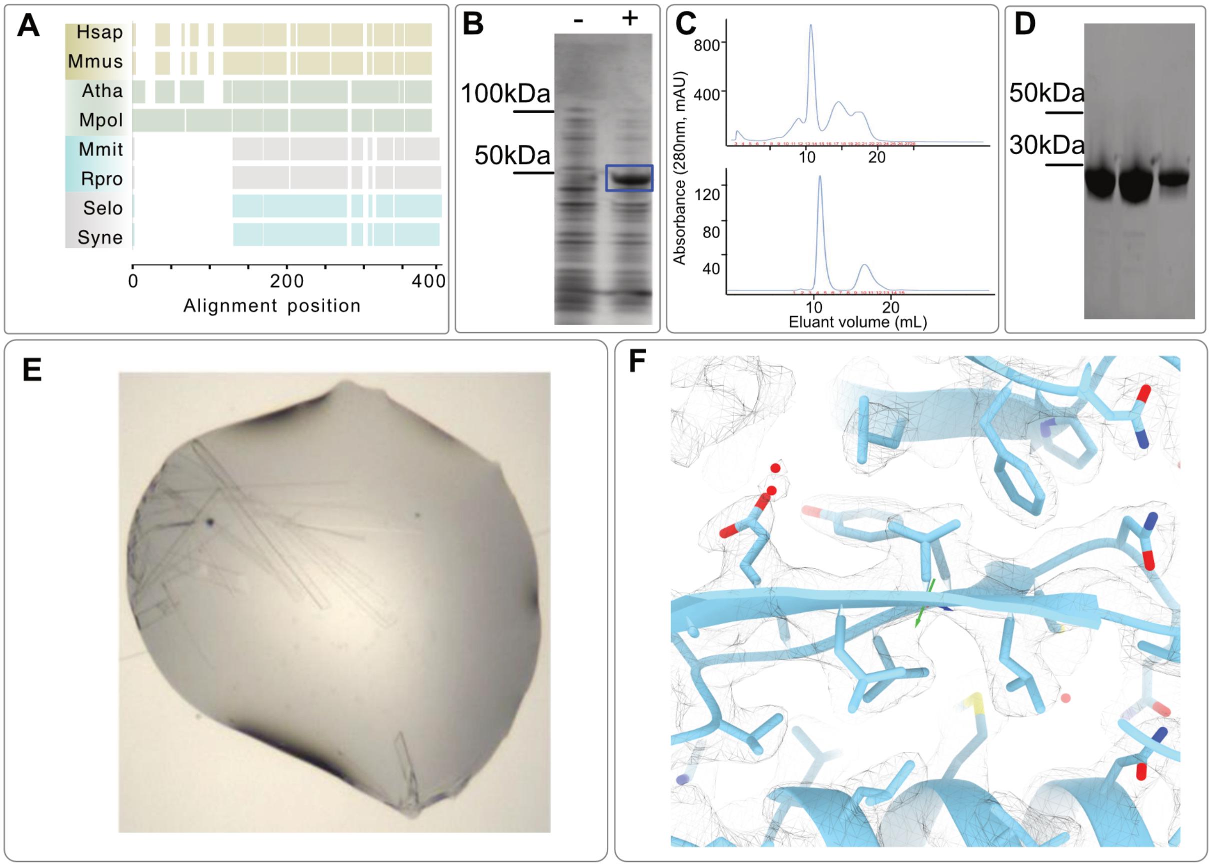

**Fig. S5: Heterologous expression, purification and crystallization of MpTACO1. (A)** Multiple sequence alignment of four eukaryotic TACO1 and four prokaryotic yebC homologs highlight the length of the targeting peptides (TP) that were added during evolution to the nuclear-encoded homologs. MpTACO1 (Mp1g23800; amino acids 127-374) N-terminally fused with 6X-Histidine and SUMO, and expressed in *E. coli* without (-) and with (+) inducer; total protein from overnight culture separated on SDS-PAGE and visualized with Coomassie Brilliant Blue (the blue box indicates expected protein band for MpTACO1). **(B-C)** Anion exchange chromatogram with major peaks in fraction 13-15 and size exclusion (Superdex™ 75 10/300 GL) chromatogram with major peaks in fractions 4-6, **(D)** which are resolved on SDS-PAGE and visualized with Coomassie Brilliant Blue. **(E)** Plates formed after 20h of crystallization in the Hcryo screen (crystallization solution: 0.1M MES, pH 6, 30% (v/v) polyethylene glycol 200, 5% (w/v) polyethylene glycol 3350). (**F)** Close up of the atomic modelling of MpTACO1 into the electron density. Density is shown as a mesh at 1 standard deviations (σ) above the mean.

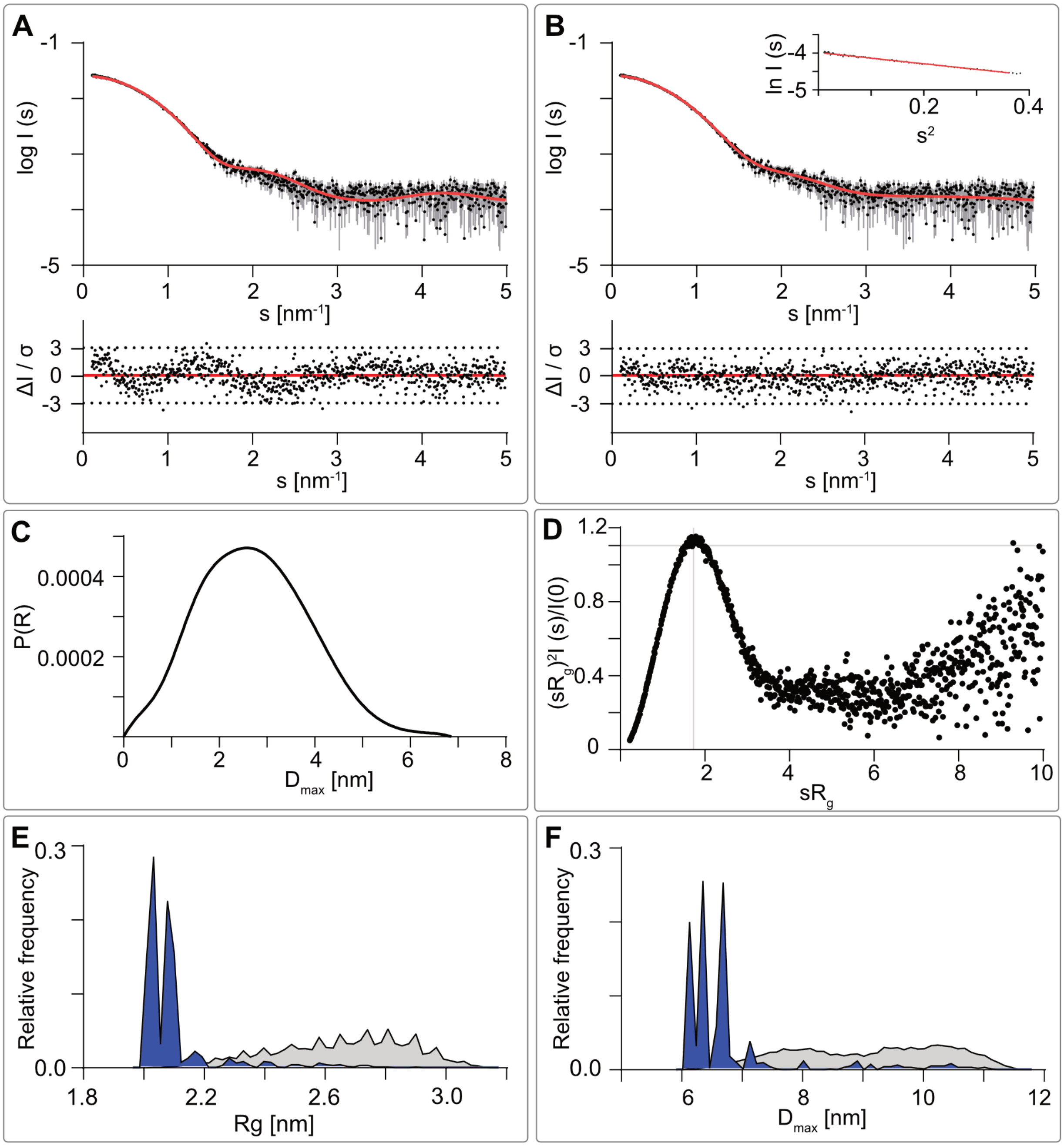

**Fig. S6: Small-angle X-ray scattering data of MpTACO1. (A)** Experimental data (black dots, with grey error bars) and the theoretical scattering intensity (red line) from the initial crystal structure calculated with CRYSOL ($\chi^2$ value 1.39). Shown below is the residual plot of the data. **(B)** Experimental data (black dots, with grey error bars) and the EOM ensemble (red line) along with the residual plot of the data at bottom. The Guinier plot is added in the top right corner. **(C)** The Distance distribution function *(p(r)* function) indicating a globular molecule with small elongation. **(D)** Dimensionless Kratky plot indicating a compact molecule with certain degree of flexibility. **(E)** $R_g$ and **(F)** $D_{max}$ distributions calculated by EOM. Random pool distribution is shown in grey and the selected models in blue.

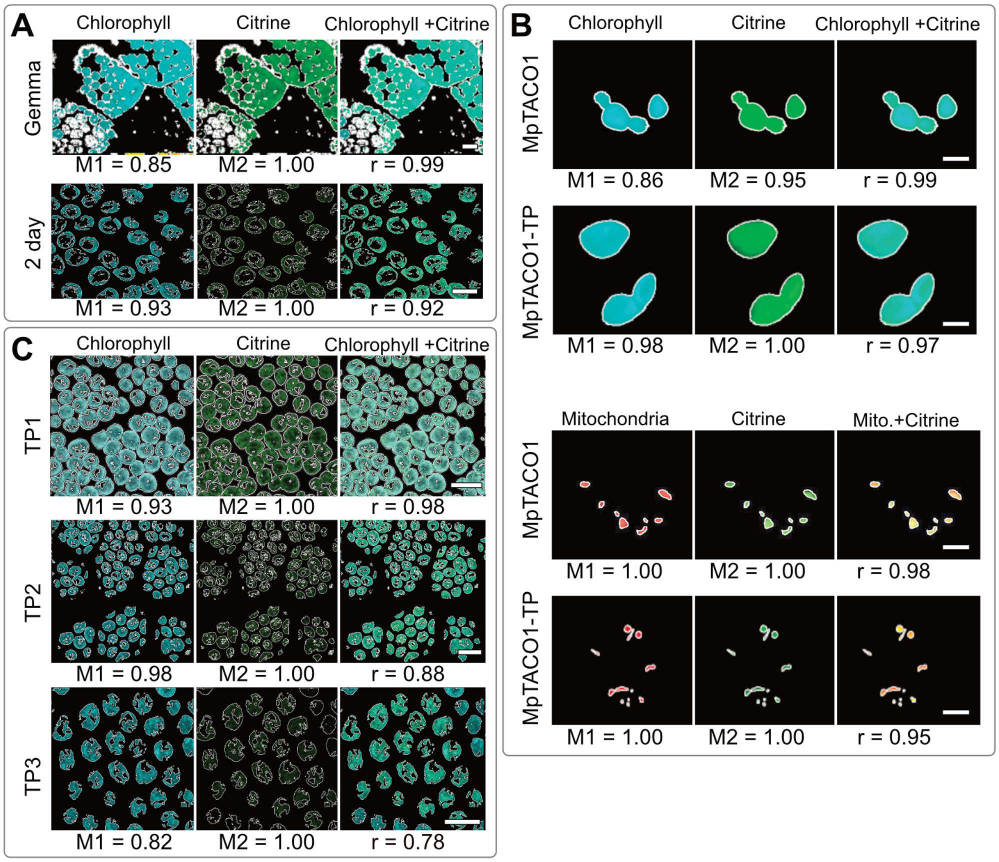

**Fig. S7: Co-localization analysis on plant lines expressing MpTACO1:Citrine, MpTACO1-TP:Citrine and variants therein.** Colocalization analyses on each image correlating the main figure 3. Mander's overlap coefficients M1 and M2, respectively, show the fraction of the organelle signal overlapping with the reporter and *vice versa*; e.g. in the first figure in S7A, M1=0.85 indicate that 85% of chlorophyll signal overlaps with GFP and M2=1 indicates that 100% of GFP overlaps with chlorophyll. R shows the correlation coefficient for intensities of the two channels. Scalebars 10µm.

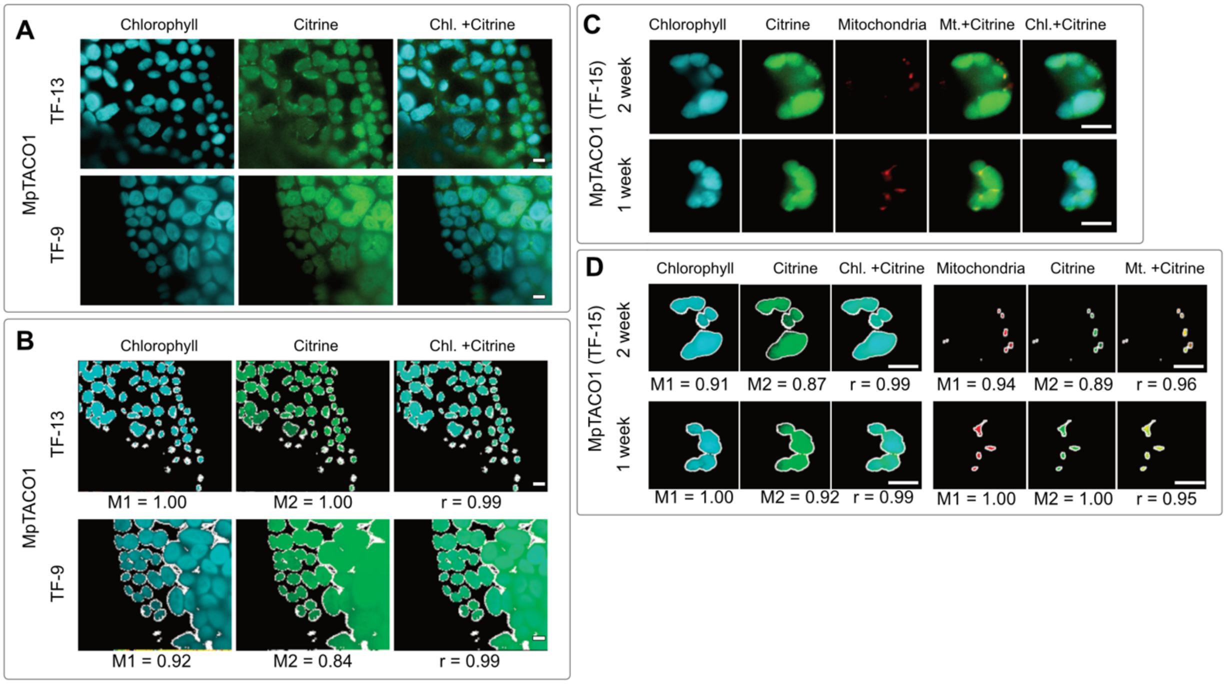

**Fig. S8: Additional microscopy images and co-localization analyses supporting the dual localization of MpTACO1. (A)** Intracellular localisation of MpTACO1 in thalli of two independent transfectant lines with the corresponding colocalization analyses in **(B)**. **(C)** Images of protoplasts generated from one- and two-week-old thalli of transfectant lines expressing full MpTACO1 with the corresponding colocalization analyses in **(D)**. Plastids were imaged through their autofluorescence and protoplast mitochondria through MitoTracker Red. Scalebars 10µm.

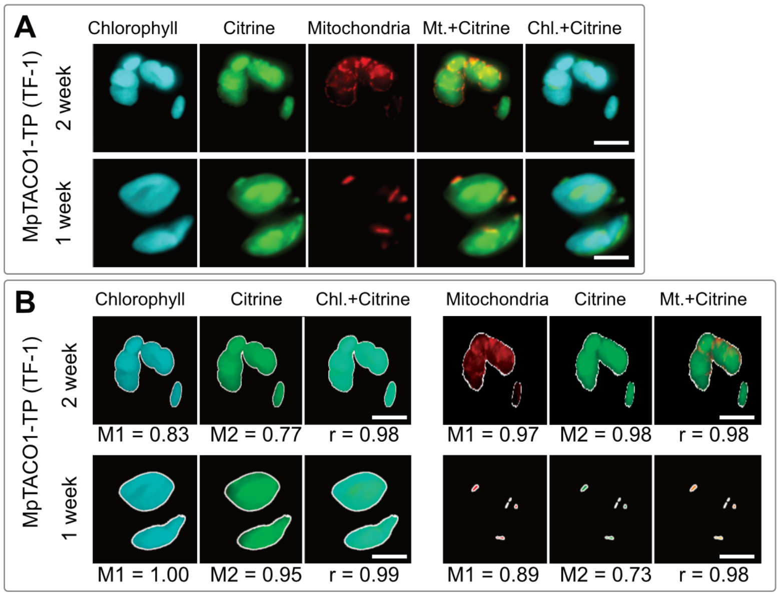

**Fig. S9: Additional microscopy images and co-localization analyses for MpTACO-TP.** Additional images of protoplast from thalli of MpTACO1-TP transfectants **(A)** and colocalization analyses **(B)**. Plastids were imaged through their autofluorescence and protoplast mitochondria through MitoTracker Red. Scalebars 10µm.

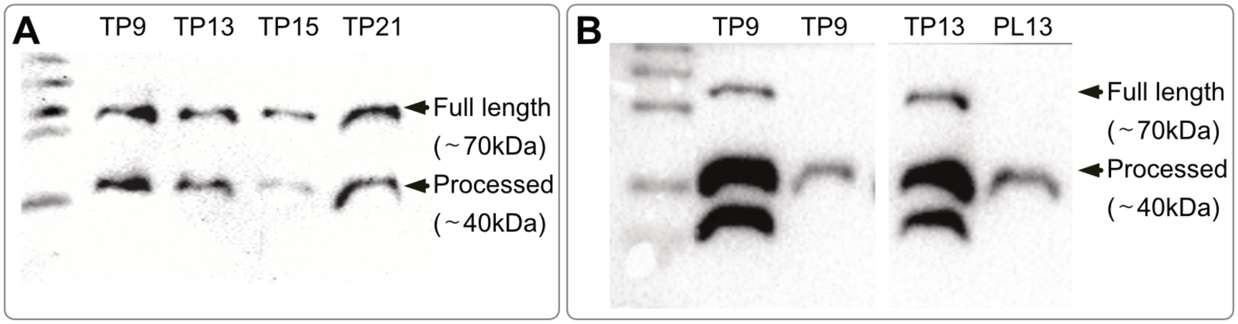

**Fig. S10: Immunoblotting from plant lines expressing MpTACO1:citrine.** Immunoblotting against citrine from total protein (TP) of four independent TACO1:citrine overexpression lines **(A)** and from total protein and isolated plastid fraction (PL) of two independent lines **(B)**.

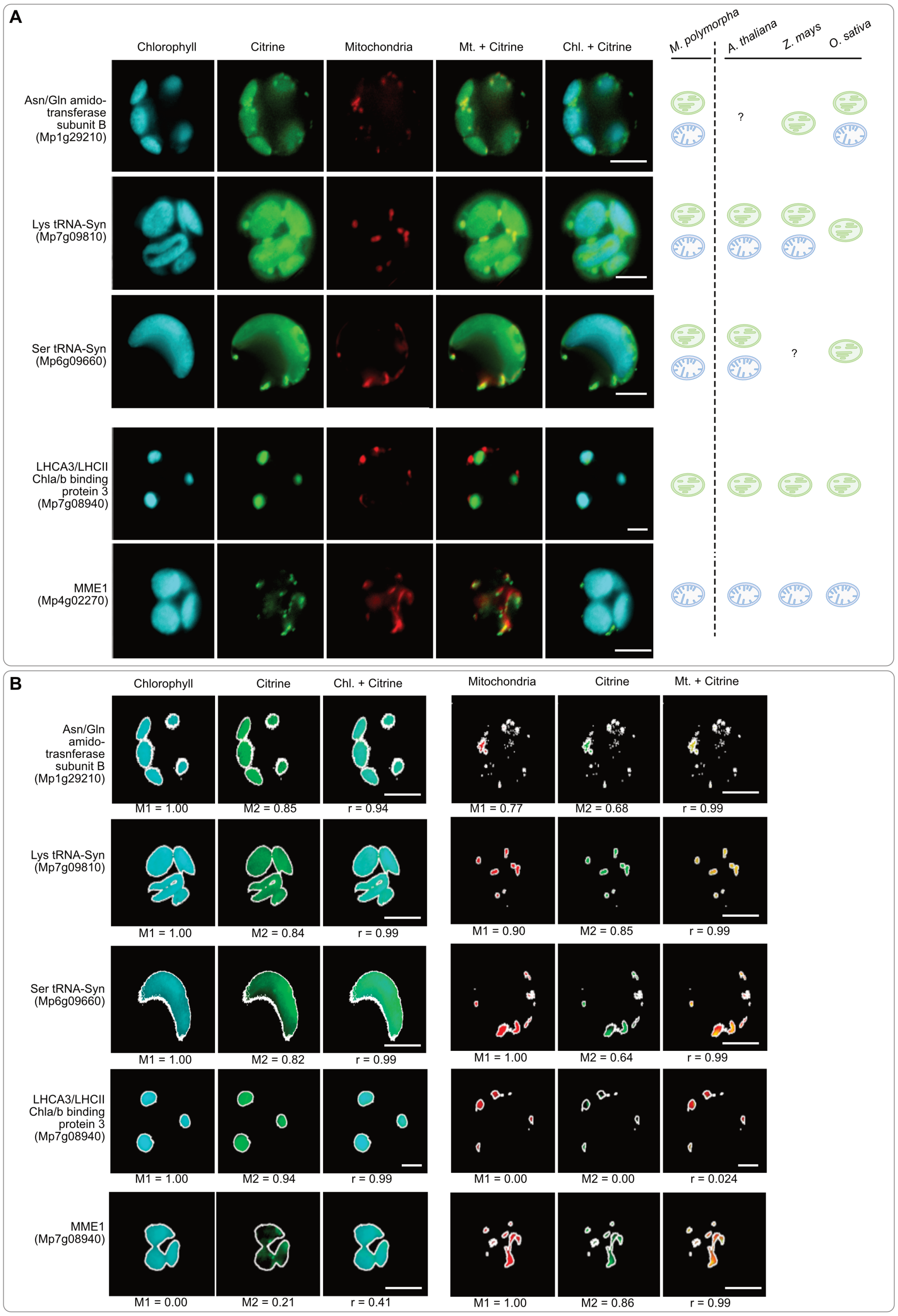

**Fig. S11: Dual targeting of three information processing-related proteins. (A)** Localization of three information processing-related candidates fused to citrine, along with plastid- and mitochondria-only controls, in protoplasts generated from thalli of one week old isogenic plant lines. Localization of homologs of these proteins, across four species, is depicted on the left. **(B)** Co-localization analyses on the same images. Plastids were imaged through their autofluorescence and protoplast mitochondria through MitoTracker Red. Scalebars 10μm.

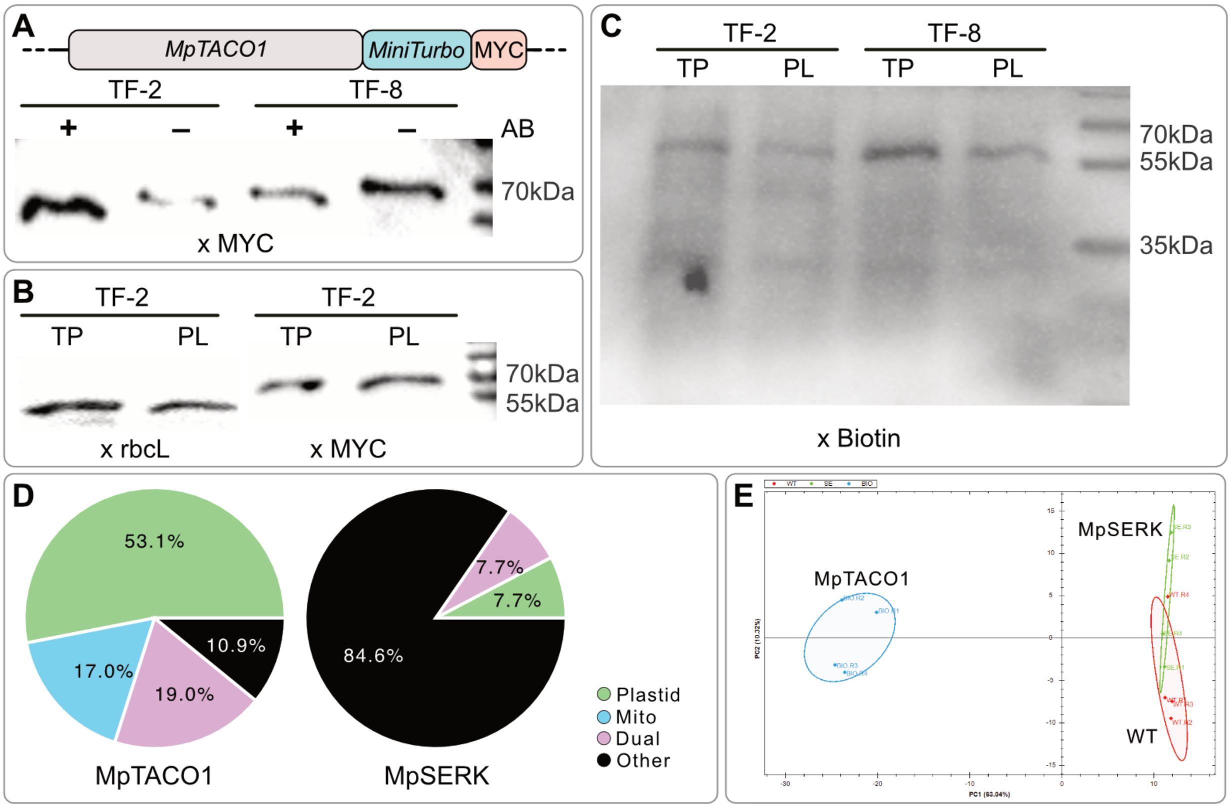

**Fig. S12: Expression and localisation of MpTACO1 fused to miniTurbo. (A)** Immunoblotting (against MYC tag) from total protein of 10-day-old thalli of two independent transfectant lines grown with (+) and without (-) antibiotic. **(B)** Immunoblotting (against rbcL and MYC tag) from total protein (TP) and plastid protein (PL) of 10-day-old thalli. **(C)** Immunoblotting (against biotin) from total protein (TP) and plastid protein (PL) of 10-day-old thalli from two independent transfectant lines, after 24h of biotin treatment. Of the two plant lines, TF-2 was used further for proximity labelling. **(D)** Distribution of the localization of proteins biotinylated by MpTACO1::miniTurbo versus the control MpSERK::miniTurbo. **(E)** Principal component analysis of the proteins enriched via mass spec in the different plant lines (n=4).

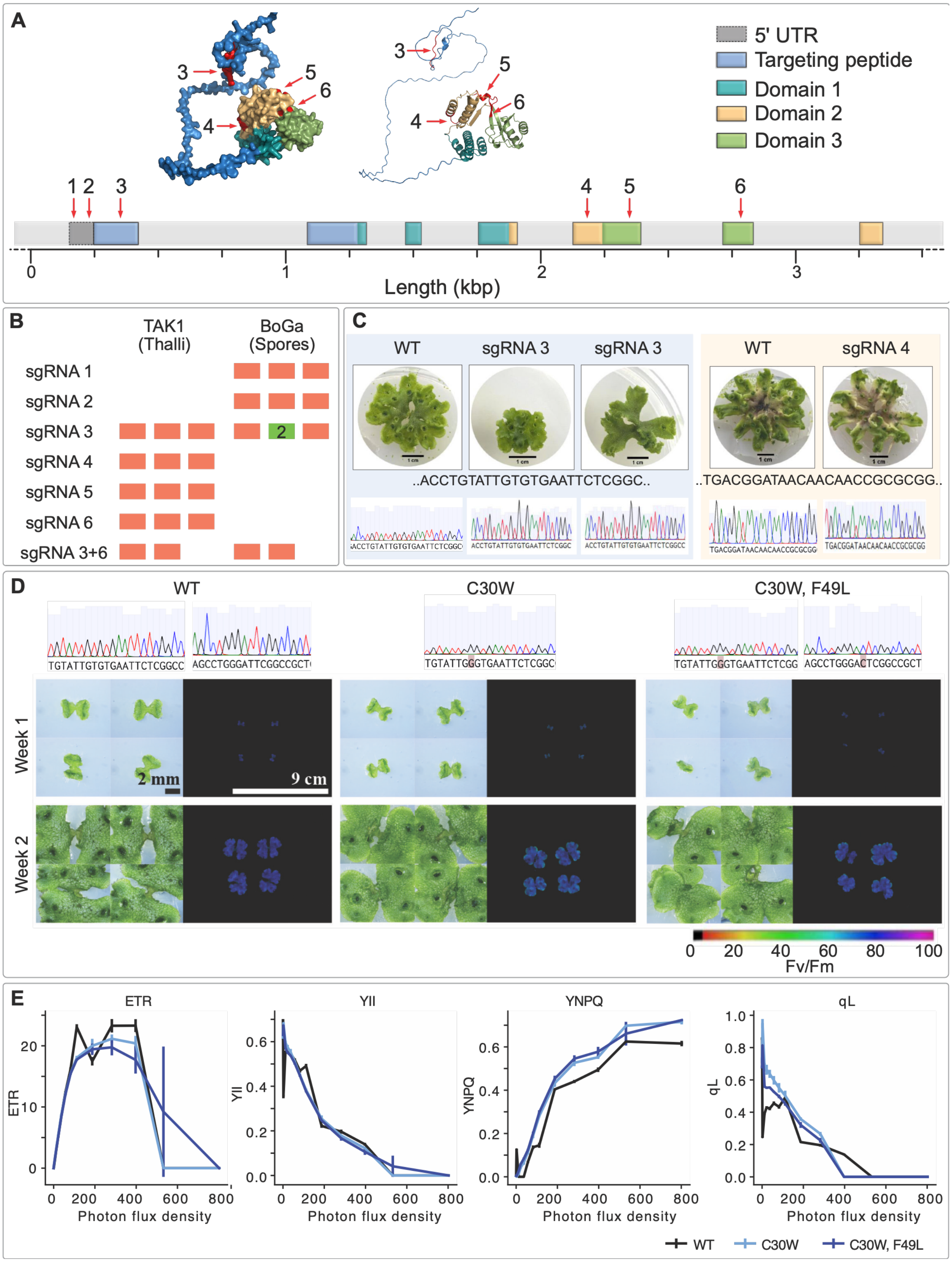
A
5' UTR
Targeting peptide
Domain 1
Domain 2
Domain 3
Length (kbp)
B
TAK1 (Thalli)
BoGa (Spores)
sgRNA 1
sgRNA 2
sgRNA 3
sgRNA 4
sgRNA 5
sgRNA 6
sgRNA 3+6
C
WT
sgRNA 3
sgRNA 3
WT
sgRNA 4
..ACCTGTATTGTGTGAATTCTCGGC..
..TGACGGATAACAACAACCGCGCGG..
D
WT
C30W
C30W, F49L
Week 1
Week 2
2 mm
9 cm
Fv/Fm
E
ETR
YII
YNPQ
qL
Photon flux density
WT
C30W
C30W, F49L

**Fig. S13: Attempts at generating loss-of-function mutants of MpTACO1 retrieve only mutant lines with silent mutations in the N-terminal targeting sequence. (A)** Four distinct regions of the coding sequence and two regions of the 5' UTR of MpTACO1 were targeted using CRISPR-Cas9 (indicated by red arrows on the gene sequence and red residues in protein's structural context). **(B)** Overview of the number of attempts to generate loss-of-function lines for each sgRNA and for two distinct strains and developmental stages; each box represents a single attempt with a minimum of 50 explants (for TAK1) or regrowing sporelings from a minimum of 25,000 spores starting culture (for BoGa). **(C)** Exemplarily, three-week-old thalli of the 4th generation after (left) and four-week-old thalli of the 4th generation (right), along with sequencing results showing no mutations in the sgRNA target region. **(D)** Sequencing results, thalli structure, and photosynthetic Fv/Fm (maximum PSII photochemical efficiency) for two plant lines with amino acid substitutions in the targeting peptide of MpTACO1. **(E)** Pulse-amplitude-modulated (PAM) chlorophyll fluorescence analysis showing apparent PSII electron transport rate (ETR), effective PSII quantum yield under actinic light (Y(II)), regulated non-photochemical energy dissipation (Y(NPQ)), and the estimated fraction of open PSII reaction centres (qL) for the two-week-old plants of (D).

**Supplementary Table S1: X-ray data collection and refinement statistics for MpTACO1**

| | |
|---|---|
| Data collection statistics | |
| Beamline | ID30A-3 |
| Space group | C 1 2 1 |
| Cell dimensions a, b, c (Å) | 91.7, 38.5, 78.1 |
| Cell dimensions α, β, γ (°) | 90, 115.6, 90 |
| Resolution range (Å) | 70.49-2.34 |
| Wavelength (Å) | 0.9677 |
| $R_{merge}$ | 0.157 (1.191)[a] |
| $R_{meas}$ | 0.184 (1.388) |
| $R_{pim}$ | 0.095 (0.710) |
| I/σ(I) | 7.4 (1.2) |
| $CC_{1/2}$ | 0.993 (0.721) |
| Completeness (%) | 97.61 (98.03) |
| Multiplicity | 7.1 (7.2) |
| Wilson B-factor (Å$^2$) | 42.65 |
| Total number of observations | 73,568 (7,298) |
| Total number unique | 10,404 (2,582) |
| Refinement statistics | |
| Resolution (Å) | 45.18-2.34 (2.57-2.34) |
| $R_{work}/R_{free}$ (%) | 23.31/27.90 |
| Reflections used for $R_{free}$ | 510 |
| Number of atoms protein/water | 3,722/28 |
| Average B-factor | 50.04 |
| R.m.s. deviations[b] | |
| Bond lengths (Å) | 0.003 |
| Bond angles (°) | 0.62 |
| Ramachandran analysis | |
| Favoured region (%) | 99.15 |
| Allowed region (%) | 0.85 |

| | |
|---|---|
| Outliers (%) | 0.0 |
| Clashscore | 1.08 |
| Rotamer outliers (%) | 0.0 |

### Supplementary Table S2: SAXS Data.

| | |
|---|---|
| **Data collection parameters** | |
| SAXS Device | Xenocs Xeuss 2.0 with Q-Xoom |
| Detector | PILATUS 3 R 300K windowless |
| Detector distance (m) | 0.55 |
| Beam size | 0.8 mm x 0.8 mm |
| Wavelength (nm) | 0.154 |
| Sample environment | Low Noise Flow Cell, 1 mm ø |
| Absolute scaling method | Comparison with scattering from pure $H_2O$ |
| Normalization | To transmitted intensity by direct beam- |
| Scattering intensity scale | Absolute scale, $cm^{-1}$ |
| *s* range ($nm^{-1}$) | 0.05 – 5.5 |
| **Sample** | **MpTACO1** |
| Organism | *Marchantia polymorpha* |
| UniProt ID | 9SCZ |
| Mode of measurement | batch |
| Temperature (°C) | 15 |
| Exposure time (# frames) | 600 s (36 frames) |
| Protein buffer | 50 mM Tris-HCl pH 8.0, 150 mM NaCl |
| Protein concentration (mg/ml) | 5.44 |
| **Structural parameters** | |
| *Guinier Analysis (PRIMUS)* | |
| $I(0) \pm \sigma$ ($cm^{-1}$) | 0.019 ± 0.00005 |
| $R_g \pm \sigma$ (nm) | 2.14 ± 0.010 |
| *s-range* ($nm^{-1}$) | 0.105 – 0.602 |
| *min < sRg < max limit* | 0.225 – 1.288 |
| Data point range | 1 - 85 |
| Linear fit assessment ($R^2$) | 0.992 |
| *PDDF/P(r) Analysis (GNOM)* | |
| $I(0) \pm \sigma$ ($cm^{-1}$) | 0.018 ± 0.00005 |
| $R_g \pm \sigma$ (nm) | 2.10 ± 0.007 |
| $D_{max}$ (nm) | 6.86 |
| Porod volume ($nm^3$) | 45.20 |
| *s-range* ($nm^{-1}$) | 0.105 – 5.356 |
| $\chi 2$ / CorMap P-value | 1.049 / 0.585 |
| **Molecular mass (kDa)** | |
| From *I*(0) | 26.31 |
| From Qp [1] | 24.07 |
| From MoW2 [2] | 27.04 |
| From Vc [3] | 28.62 |
| From Bayesian Inference [4] | 26.23 |
| From GNNOM [5] | 27.30 |
| From sequence | 26.98 (monomer) |
| **Atomistic modelling** | |
| EOM | |

| | |
|---|---|
| Symmetry | P1 |
| *s*-range for fit ($nm^{-1}$) | 0.105 – 4.988 |
| $\chi^2$, CorMap *P*-value | 1.024 / 0.808 |
| **SASBDB accession codes** [6] | SASDYA7 |
| **Software** | |
| ATSAS Software Version [7] | 3.0.5 (ATSAS online 3.2.1) |
| Primary data reduction | PRIMUS [8] |
| Data processing | GNOM [9] |
| Flexibility ensemble modelling | EOM [10,11] |
| Statistic goodness-of-fit test | $\chi^2$, CorMap [12] |
